# Accurate assignment of disease liability to genetic variants using only population data

**DOI:** 10.1101/2021.04.19.440463

**Authors:** Joseph M. Collaco, Karen S. Raraigh, Joshua Betz, Melis Atalar Aksit, Nenad Blau, Jordan Brown, Harry C. Dietz, Gretchen MacCarrick, Lawrence M. Nogee, Molly B. Sheridan, Hilary J. Vernon, Terri H. Beaty, Thomas A. Louis, Garry R. Cutting

## Abstract

**Purpose:** The growing size of public variant repositories prompted us to test the accuracy of predicting pathogenicity of DNA variants using population data alone.

**Methods:** Under the *a priori* assumption that the ratio of the prevalence of variants in healthy and affected populations form two distinct distributions (pathogenic and benign), we used a Bayesian method to assign probability of a variant belonging to either distribution.

**Results:** The approach, termed BayPR, accurately parsed 300 of 313 expertly curated *cystic fibrosis transmembrane conductance regulator (CFTR)* variants: 284 of 296 pathogenic/likely pathogenic (P/LP) variants in one distribution and 16 of 17 benign/likely benign (B/LB) variants in another. BayPR produced an area under the receiver operating curve (AUC) of 0.99 for 103 functionally-confirmed missense *CFTR* variants, equal to or exceeding ten commonly used algorithms (AUC range: 0.54 to 0.99). Application of BayPR to expertly curated variants in eight genes associated with seven Mendelian conditions assigned ≥80% disease-causing probability to 1,350 of 1,374 (98.3%) P/LP variants and ≤20% to 22 of 23 (95.7%) B/LB variants.

**Conclusion:** Agnostic to variant type or functional effect, BayPR provides probabilities of pathogenicity for DNA variants responsible for Mendelian disorders using *only* variant counts in affected and unaffected population samples.

## INTRODUCTION

In an era of rapidly expanding genetic testing, there is a growing challenge for clinical and research laboratories to determine the phenotypic significance and clinical consequences of DNA variants. Many algorithms predict disease liability using population data, predictive tools, segregation analysis, novelty, allelic information, and prior reporting in expertly curated locus specific or confederated databases^1-3^. The overall trend has been to combine an ever-increasing number of tools; the most accurate are amalgams of existing prediction algorithms^4,5^. This approach is fraught with the problem of circularity, as many of the incorporated tools use the same variant features and training sets^6^. Indeed, it is unclear whether continued development of ever more complex algorithms will achieve greater accuracy. We therefore sought to determine if we could improve the predictive potential of one individual data element used to predict pathogenicity.

The widely adopted standards and guidelines devised by the American College of Medical Genetic (ACMG)/Association of Molecular Pathology (AMP) integrate population data as part of their criteria^7^. The primary assumption when including population data is that the frequency of a pathogenic variant in a collection of affected individuals should exceed that observed in a sample of individuals without the disease of interest. However, the ACMG/AMP recommended comparison of variant frequencies in affected and unaffected samples using odds ratios with a somewhat arbitrary threshold is problematic when applied to Mendelian disorders with different penetrance characteristics. Furthermore, the population criterion becomes less effective as variant frequency decreases and prevalence cannot be meaningfully compared for variants that may be observed once or only a few times in a sample of diagnosed individuals, or not observed in a large sample of healthy and presumed unaffected individuals. Finally, the categorical nature of assigning this information to different levels of confidence regarding pathogenicity does not accommodate the spectrum of phenotypic consequences conferred by allelic heterogeneity^8^.

Bayesian methods are well-suited to address this problem, as they can incorporate existing sources of information to infer the likelihood of a very rare or even unobserved event^9^. Sources may include orthogonal data from the subjects, prior probabilities informed by previous studies, or pooled information across individuals. Consequently, we developed and tested a population-based approach within a Bayesian framework, termed Bayesian Prevalence Ratio or BayPR. Using only population data, namely the prevalence of variants in samples of diagnosed and unaffected individuals, we demonstrate that most variants fall into one of two distributions, under the *a priori* assumption that pathogenic or benign variants would self-sort into separate groups^9^. Critically, the sorting would occur agnostic of functional data or other metrics of disease liability. We demonstrate the utility of this approach using the Genome Aggregation Database (gnomAD)^10^ as a ‘healthy’ population sample, and a worldwide collection of individuals diagnosed with CF [MIM: 219700] (caused by variation in *CFTR*) as the ‘disease’ sample^11^. We apply BayPR to expertly curated variants associated with Mendelian disorders with different inheritance patterns to demonstrate the generalizability of our approach.

## SUBJECTS AND METHODS

### Data Sources

All variants were de-identified. <u>**“Control” dataset**</u>: Aggregated genomic data were downloaded from gnomAD v2.1.1, a publicly available database consisting of 125,748 exome sequences and 15,708 whole genome sequences from unrelated individuals^10^. <u>**Cystic Fibrosis (*CFTR*):**</u> *CFTR* genotype data were provided by the CFTR2 [<u>C</u>linical and <u>F</u>unctional <u>TR</u>anslation of CFTR] (https://cftr2.org) database^11^. <u>**Phenylketonuria (*PAH*):**</u> *PAH* genotype data were provided by the BIOPKU database^12^. <u>**Interstitial Lung Disease (*ABCA3*):**</u> Variants were assembled from individuals with genetic surfactant dysfunction studied in research laboratories at Washington University School of Medicine and Johns Hopkins University School of Medicine and from publications^13^. <u>**X-linked Adrenoleukodystrophy (*ABCD1*):**</u> *ABCD1* variants were obtained from clinical testing in the Johns Hopkins Genomics DNA Diagnostic lab from 2016 to 2020. <u>**Barth Syndrome (TAZ):**</u> *TAZ* variants were obtained from the Human Tafazzin Gene Variants Database (a sub-database of the Barth Syndrome Registry and Repository; https://www.barthsyndrome.org/research/tafazzindatabase.html)^14^. <u>**Marfan Syndrome (*FBN1*):**</u> Variants were identified during clinical testing at various CLIA-approved commercial labs from 2006 to 2020. <u>**Loeys-Dietz Syndrome (*TGFBR1/TGFBR2*):**</u> Variants were identified during clinical testing at various CLIA-approved commercial labs from 2006 to 2020. Further information concerning curation for all populations can be found in the **supplementary material**. All genetic data (control and disease populations) and disease liability assignments for alleles can be found in the **data supplement**.

### Analytic method

An empirical Bayesian approach was used to analyze the counts of variants in the disease-specific population database relative to the control gnomAD population using a two component finite mixture model. Each component modeled the number of variants arising from cases conditional on the total number of observed variants, using a binomial likelihood, placing a beta prior on a transformation of the prevalence ratio (see **supplementary material** for detailed information). This approach converts the proportion of variants observed in cases to the prevalence ratio in cases relative to controls. Estimates were obtained by maximizing the marginal likelihood using the expectation-maximization (EM) algorithm^15^, implemented using the optimx package in R (R Foundation for Statistical Computing, Vienna, Austria)^16^. A grid search was used to assess sensitivity to starting values for the EM algorithm, including the parameters of each mixture component and the mixing fraction. The model allows for variation in the number of individuals assessed for a given variant in each database but does assume a relative constant ratio of individuals in the affected database versus the control database across all variants; variation from this ratio would induce a different prior on the prevalence ratio. The R program used to generate the probabilities can be found on github (https://github.com/melishg/BayPR/). The BayPR method was compared to other predictive algorithms using non-parametric estimations of ROC curves with Bamber and Hanley confidence intervals for the AUCs; significance levels were adjusted for via Sidak’s correction. Heatmaps were generated using the contour command in STATA IC 15, which uses thin plate splines for interpolation. For sensitivity and specificity analyses, score thresholds were set to determine whether a variant had a positive or negative result from the BayPR. For P/LP variants, BayPR scores >90% or >80% were considered true positives. For B/LB variants, Bayesian scores <10% or <20% were considered true negatives. Determinations of P/LP and B/LB status were made by expert review. Only genes with both P/LP and B/LB variants underwent sensitivity and specificity analysis.

## RESULTS

### Predicting disease liability of *CFTR* variants using prevalence ratios

Our underlying premise is based on the differences in prevalence of pathogenic and benign variants among ‘healthy’ and ‘affected’ samples. For example, plotting 22 CFTR variants (all established as pathogenic for cystic fibrosis (CF) by an ACMG expert committee in 2004^17^) as a ratio of their prevalence in affected and unaffected samples reveals clear deviation from neutrality (**Figure 1A**). Each variant individually generates a relative risk greater than 5.0 with confidence intervals that do not include 1.0, thereby satisfying ACMG/AMP criteria for pathogenicity^7^. However, application of relative risk or odds ratios is problematic for many rare pathogenic variants and cannot be used to evaluate benign variants. In contrast, prevalence ratios cluster 152 common and rare expertly curated CFTR variants into distinct pathogenic/likely pathogenic (P/LP) and benign/likely benign (B/LB) groups (**Figure 1B**)^8,11,18^. To derive the probability of a variant being in one group or the other, we implemented a two-component finite mixture model in an empirical Bayesian framework by pooling observed information across the different variants^15^ (see **Supplementary Methods** for detailed explanation). The model generates 95% credible intervals for the prevalence ratios based on observed allele counts and then groups variants according to their similarity to other variants with regards to observed frequencies. The combining of information generates better estimates of each individual variant prevalence ratio and allows assessment of the probability that a variant came from the population with higher- or lower-than-average prevalence ratios. The posterior probability derived from a prevalence ratio estimates the likelihood that any variant belongs in one of two distributions. We first applied our method, termed Bayesian Prevalence Ratio or BayPR, to 313 *CFTR* variants interpreted using clinical, functional, and segregation information (https://cftr2.org)^8,11,18^. Parsing of variants based on probabilities using BayPR placed variants into two distinct distributions: 284 out of the 296 P/LP variants in one distribution using probability threshold of >90%, while 16 out of the 17 B/LB variants are in the second distribution based on having <10% probability of being assigned to the first distribution (**Figure 1C and Table 1**). The probability of a variant being assigned to one distribution or the other followed expectations for mechanism. For example, 185 of 188 (98.4%) null variants (e.g., splice donor/acceptor, frameshift, and nonsense) that can be confidently interpreted as P/LP using ACMG/AMP variant guidelines^7^ had probabilities placing them in the ‘disease’ distribution (**Figure 1D**). Variants that may allow residual CFTR function (e.g., missense and synonymous) appeared in both distributions, primarily in accordance with disease liability established from functional studies^8^. BayPR generated an area under the ROC curve (AUC) of 0.99 (95%CI: 0.98, 1.00) for 101 expertly interpreted missense CFTR variants (**Figure 1E**). This was matched only by ClinPred^19^ (0.99; 95%CI: 0.97, 1.00), which is remarkable given that *BayPR is based only on population data*, while ClinPred uses allele frequency data, prediction scores from 16 algorithms, and is trained on variants from the ClinVar database^20^. The AUCs of other commonly utilized predictors ranged from 0.54 to 0.87 (Bonferroni corrected *p* values compared to BayPR ranged from <0.001 to 0.03). To assess the utility of BayPR for low frequency variants, we generated probabilities for 161 P/LP variants in CFTR2 having counts of 20 or fewer. Of these, BayPR assigned >90% probability to 153 (95%) indicating that the method performs equally well on lower-frequency variants as it does on more frequent variants. This attribute will be useful in the clinical situation when a rare variant is identified. As an example, if a diagnostic genetic lab were to identify a single novel *CFTR* variant not observed in gnomAD, BayPR generates a probability of 96.3%, placing it in the disease distribution; if the same variant were observed as few as three times in gnomAD, then then BayPR probability of belonging to the disease distribution drops to 6.8% **(Figure 1F)**.

**Table 1.**
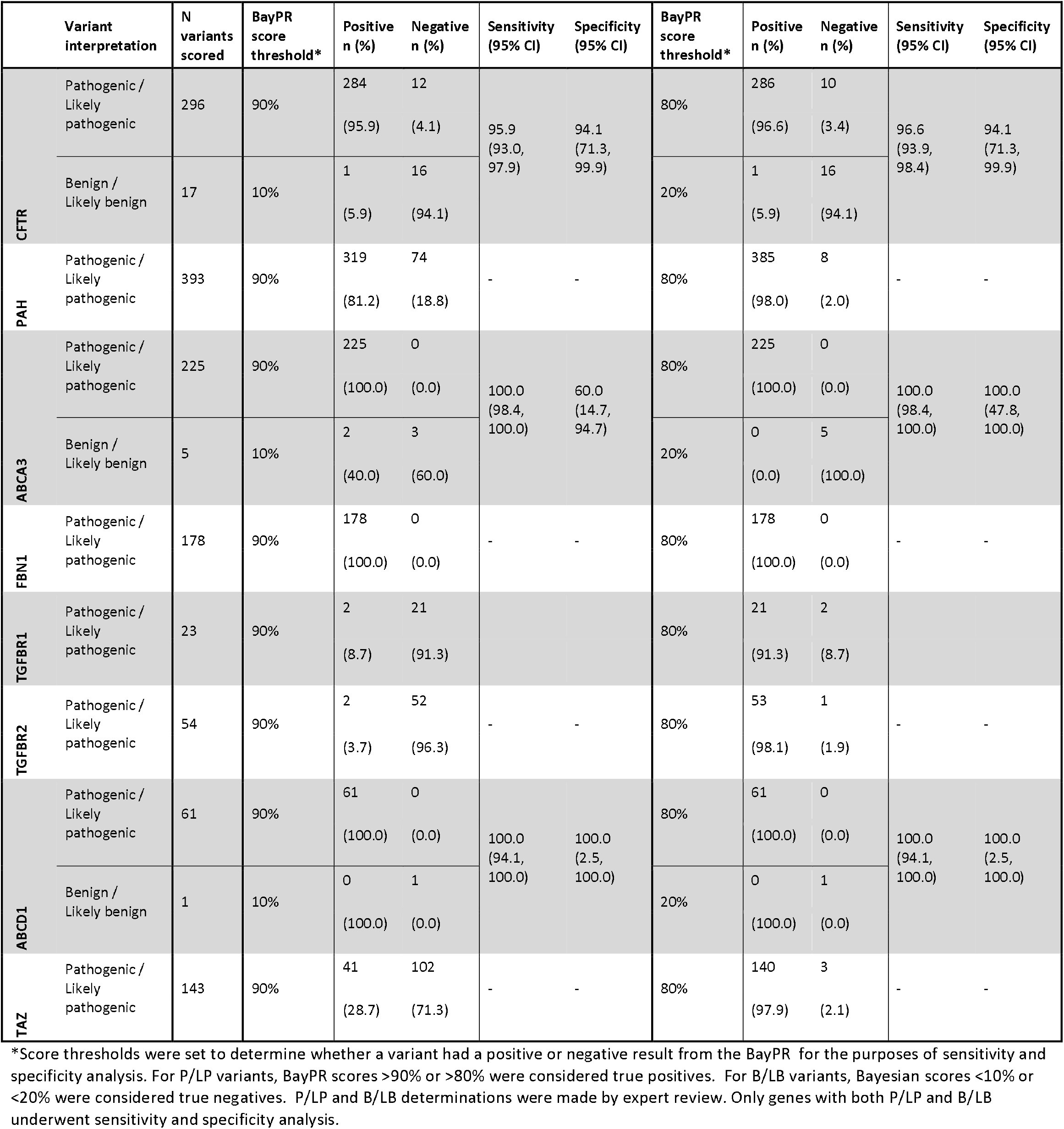
Sensitivity and specificity analysis of BayPR.

|  | Variant interpretation | N variants scored | BayPR score threshold* | Positive n (%) | Negative n (%) | Sensitivity (95% CI) | Specificity (95% CI) | BayPR score threshold* | Positive n (%) | Negative n (%) | Sensitivity (95% CI) | Specificity (95% CI) |
| --- | --- | --- | --- | --- | --- | --- | --- | --- | --- | --- | --- | --- |
| CFTR | Pathogenic / Likely pathogenic | 296 | 90% | 284<br>(95.9) | 12<br>(4.1) | 95.9<br>(93.0, 97.9) | 94.1<br>(71.3, 99.9) | 80% | 286<br>(96.6) | 10<br>(3.4) | 96.6<br>(93.9, 98.4) | 94.1<br>(71.3, 99.9) |
|  | Benign / Likely benign | 17 | 10% | 1<br>(5.9) | 16<br>(94.1) |  |  | 20% | 1<br>(5.9) | 16<br>(94.1) |  |  |
| PAH | Pathogenic / Likely pathogenic | 393 | 90% | 319<br>(81.2) | 74<br>(18.8) | - | - | 80% | 385<br>(98.0) | 8<br>(2.0) | - | - |
| ABCA3 | Pathogenic / Likely pathogenic | 225 | 90% | 225<br>(100.0) | 0<br>(0.0) | 100.0<br>(98.4, 100.0) | 60.0<br>(14.7, 94.7) | 80% | 225<br>(100.0) | 0<br>(0.0) | 100.0<br>(98.4, 100.0) | 100.0<br>(47.8, 100.0) |
|  | Benign / Likely benign | 5 | 10% | 2<br>(40.0) | 3<br>(60.0) |  |  | 20% | 0<br>(0.0) | 5<br>(100.0) |  |  |
| FBN1 | Pathogenic / Likely pathogenic | 178 | 90% | 178<br>(100.0) | 0<br>(0.0) | - | - | 80% | 178<br>(100.0) | 0<br>(0.0) | - | - |
| TGFBR1 | Pathogenic / Likely pathogenic | 23 | 90% | 2<br>(8.7) | 21<br>(91.3) | - | - | 80% | 21<br>(91.3) | 2<br>(8.7) | - | - |
| TGFBR2 | Pathogenic / Likely pathogenic | 54 | 90% | 2<br>(3.7) | 52<br>(96.3) | - | - | 80% | 53<br>(98.1) | 1<br>(1.9) | - | - |
| ABCD1 | Pathogenic / Likely pathogenic | 61 | 90% | 61<br>(100.0) | 0<br>(0.0) | 100.0<br>(94.1, 100.0) | 100.0<br>(2.5, 100.0) | 80% | 61<br>(100.0) | 0<br>(0.0) | 100.0<br>(94.1, 100.0) | 100.0<br>(2.5, 100.0) |
|  | Benign / Likely benign | 1 | 10% | 0<br>(100.0) | 1<br>(0.0) |  |  | 20% | 0<br>(100.0) | 1<br>(0.0) |  |  |
| TAZ | Pathogenic / Likely pathogenic | 143 | 90% | 41<br>(28.7) | 102<br>(71.3) | - | - | 80% | 140<br>(97.9) | 3<br>(2.1) | - | - |

\*Score thresholds were set to determine whether a variant had a positive or negative result from the BayPR for the purposes of sensitivity and specificity analysis. For P/LP variants, BayPR scores >90% or >80% were considered true positives. For B/LB variants, Bayesian scores <10% or <20% were considered true negatives. P/LP and B/LB determinations were made by expert review. Only genes with both P/LP and B/LB underwent sensitivity and specificity analysis.

**Figure 1.**
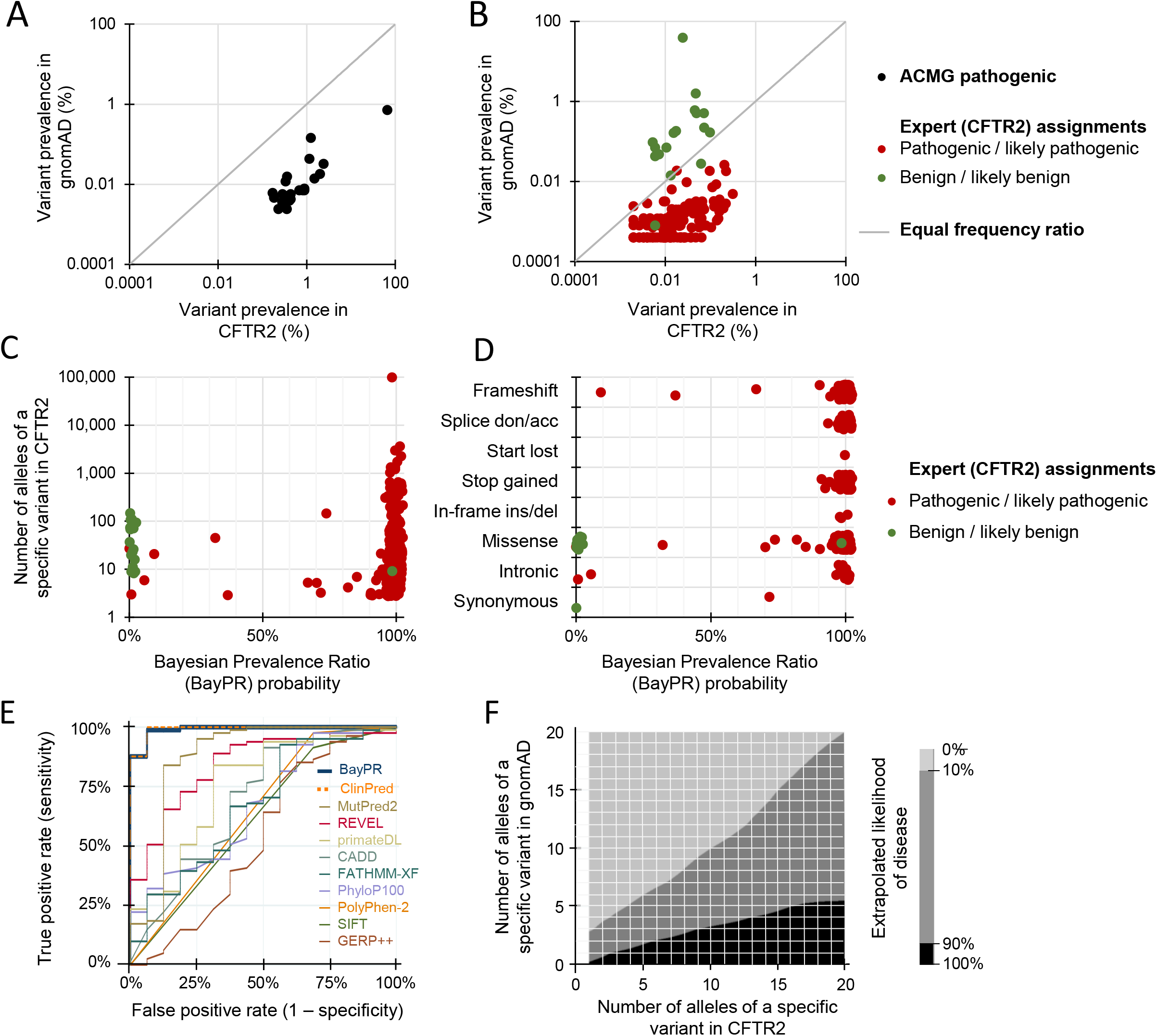
Differentiating CFTR variants using Bayesian Prevalence Ratios (BayPR). Data is jittered for display purposes only in **Figures 1A-D**; precise allele counts and BayPR probabilities are provided in the Data Supplement. **A** Plot of the prevalence ratios (PR) of 22 of the most common pathogenic *CFTR* variants (black dots) according frequencies in affected (CFTR2) and healthy (gnomAD) individuals demonstrates expected deviation from neutrality (grey line). The designation of each of these variants as pathogenic by the American College of Medical Genetics (ACMG) remains unchanged since their initial interpretation in 2004. **B** Plot of the prevalence ratios of 135 pathogenic or likely pathogenic (P/LP; red dots) and 17 benign or likely benign (B/LB; green dots) variants interpreted by CFTR2 that have been observed at least once in gnomAD reveals two relatively distinct populations. **C** Plot of *CFTR* variants according allele count and BayPR probability of being disease-causing. All 313 variants have been interpreted by CFTR2 as CF-causing (pathogenic/likely pathogenic [P/LP]; n=296) or not CF-causing (benign/likely benign [B/LB]; n=17) using clinical and functional criteria. **D** BayPR probabilities of 313 *CFTR* variants stratified according to their functional consequence. **E** Receiver operating characteristic (ROC) curves were plotted for BayPR and ten in silico prediction algorithms using data from 101 *CFTR* missense variants. The reference standard are annotations in the CFTR2 database based on clinical and in vitro data. **F** Heatmap depicts estimated probability of a *CFTR* variant being associated with disease based on absolute counts of variants in the CFTR2 and gnomAD databases. The area in black represents a greater than 90% probability of being associated with CF and the area in light grey less than 10% probability. Importantly, this heatmap is only applicable to the CF data presented in this manuscript and does not apply to other genetic disorders. This heatmap was generated using the contour command in STATA IC 15, which uses thin plate splines for interpolation.

### Predicting disease liability of unassigned *CFTR* variants

The status of 969 very rare/private *CFTR* variants had not been assigned at the time of this analysis (i.e., variants of unknown significance or VUS). Application of BayPR assigned probabilities exceeding 90% for 735 (75.9%) of these 969 VUS, suggesting they are likely to be disease-causing (i.e., P/LP) while 129 (13.3%) had probabilities less than 10%, indicating they are most likely not disease-causing (i.e., B/LB; **Figure 2A**). To independently evaluate these BayPR assignments, we sorted according to predicted functional effect. BayPR correctly assigned P/LP probabilities to 378 of the 404 (93.6%) null variants (**Figure 2B**). To assess variants of varying functional effects (missense, etc), 52 randomly selected variants were independently evaluated by the CFTR2 committee using the same criteria applied to all prior annotated variants^11^. Of the 52, 35 of 37 variants interpreted as P/LP had probabilities exceeding 90% and all 3 variants annotated as B/LB had 0% probability (**Figure 2C**). Stratification of newly interpreted variants by predicted functional consequence correlated with probability; 19 of 20 null variants and 16 of 17 missense variants newly-assigned as P/LP had BayPR >90% (**Figure 2D**). We then compared ClinVar assignments of 41 variants that had not been expertly curated but had “CF” listed as a clinical phenotype, multiple submitters with no conflicts (2-star rating or higher) and disease annotation of P/LP or B/LB (**Figure 2E**). Twenty-eight of 34 variants with P/LP designations in ClinVar had probabilities >90% and all 7 B/LB variants had probabilities <10%. Stratification by predicted functional effect were consistent with BayPR probabilities for the majority of variants (**Figure 2F**). Notably, the four null variants that had probabilities less than 90% (622-1G->A [1.8%], 2215insG [54.0%], 3500-2A->T [72.0%], and 3849+1G->A [81.9%]) occur in populations under-represented in CFTR2.

**Figure 2.**
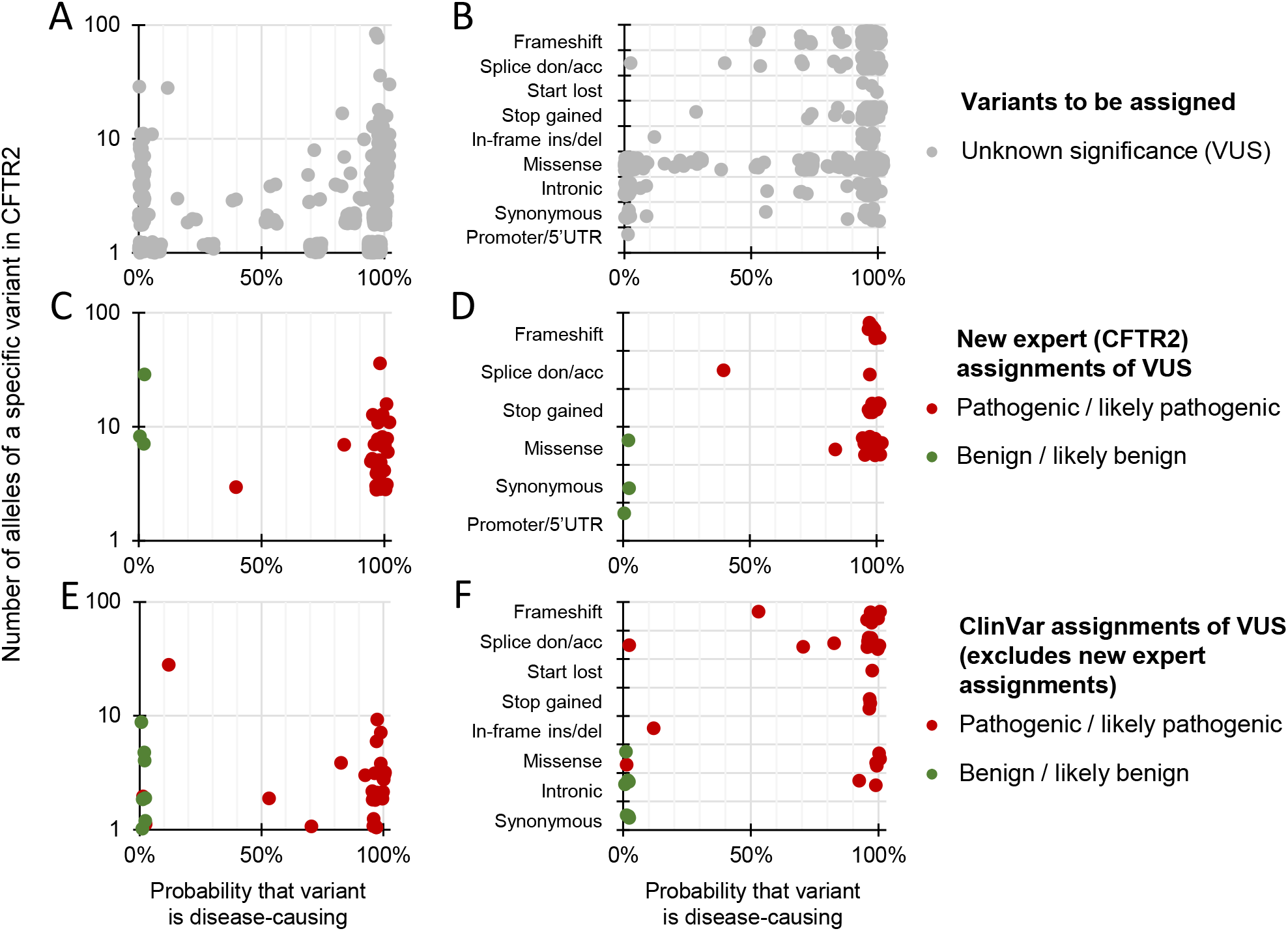
Differentiating CFTR variants of unknown significance with BayPR. Data is jittered for display purposes only in **Figures 2A-F**; precise allele counts and BayPR probabilities are provided in the Data Supplement. **A** CFTR variants that had not undergone interpretation by CFTR2 (n = 969; grey dots) cluster primarily into high (>90%; 735 variants) or low (<10%; 129 variants) probabilities. **B** BayPR probabilities of unassigned CFTR variants stratified according to their functional consequence. **C** Fifty-two unassigned CFTR variants were subsequently and independently assessed for disease liability by CFTR2 using clinical and functional criteria. **D** BayPR probabilities of newly-assigned CFTR variants stratified according to their functional consequence. **E** CFTR variants that remain without a CFTR2 interpretation and which have P/LP or B/LB interpretations in ClinVar (n = 41) cluster primarily into high (>90%; 28 variants) or low (<10%; 9 variants) probabilities. **F** BayPR probabilities of ClinVar-assigned P/LP or B/LB CFTR variants without a CFTR2 annotation parsed by predicted functional effect.

### Predicting disease liability of variants associated with other Mendelian disorders using BayPR

Having established the utility of BayPR for variants in *CFTR*, we applied this method to two other autosomal recessive disorders: phenylketonuria (PKU) caused by defects in phenylalanine hydroxylase (*PAH*; estimated 1:23,930 live births globally^12^) and interstitial lung disease (ILD) caused by dysfunction of ATP-binding cassette sub-family A member 3 (*ABCA3*; estimated 1:3,100 in Caucasian to 1:18,000 African-American live births in the U.S.)^21^. We determined BayPR scores for 726 *PAH* variants that had been evaluated by an expert panel (BIOPKU)^12^. The BIOPKU collection differed from CFTR2 in that none of the *PAH* variants had been assigned as B/LB. Despite the absence of known ‘negatives,’ BayPR assigned high disease probabilities to almost all P/LP variants. Of 334 *PAH* variants associated with classic PKU, 329 had probabilities >80%, as did 56 of the 59 mild PKU-causing variants (**Figure 3A, left panel**). Stratifying variants according to predicted function further demonstrated disease assignments were consistent, given that 169 of 174 missense variants assigned as P/LP had probabilities above 80% (**Figure 3A, right panel**). Variants in *ABCA3* were expertly interpreted using ACMG/AMP criteria^7^ (225 P/LP; 5 B/LB; and 102 VUS). The *ABCA3* variants distributed into two distinct groups: all 225 P/LP variants had probabilities >90% while all 5 B/LB variants clustered in the non-disease-causing component (**Figure 3B, left panel**). When sorted according to predicted effect, all 75 P/LP null variants and all 135 missense variants assigned as P/LP had probabilities >90% (**Figure 3B, right panel**).

**Figure 3.**
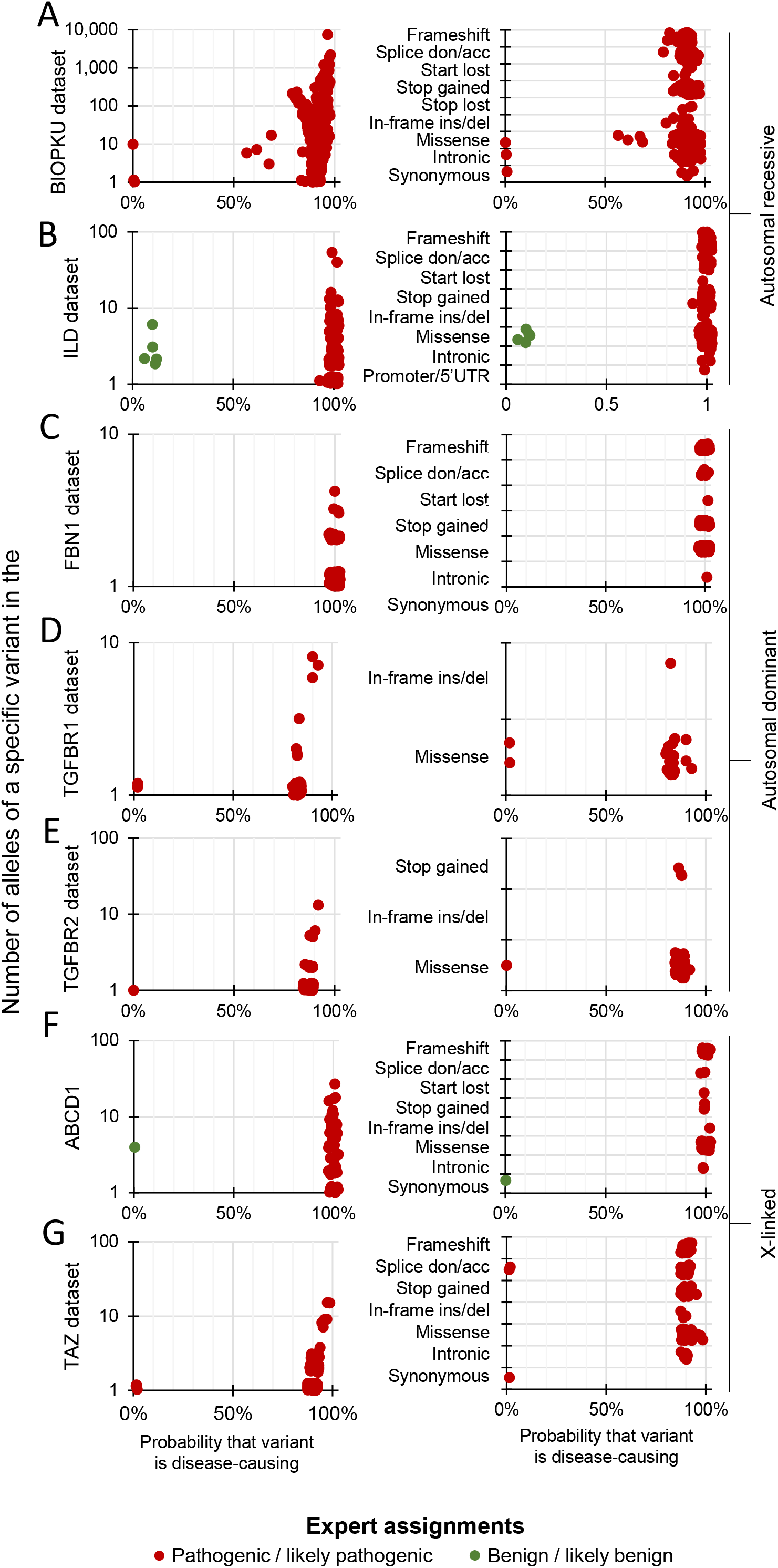
Differentiating expert assigned variants in genes associated with recessive, dominant and X-linked disorders using BayPR. Data is jittered for display purposes only in **Figures 3A-G**; precise allele counts and BayPR probabilities are provided in the Data Supplement. **A, left panel** 393 PAH variants are plotted by allele count and BayPR probabilities, with disease assignment of either pathogenic or likely pathogenic (P/LP; red dots). **A, right panel** Parsing PAH variants by predicted functional effect. **B, left panel** 230 ABCA3 (autosomal recessive interstitial lung disease) variants plotted by allele counts and probabilities, with disease assignments of P/LP (red dots) or benign/likely benign (B/LB; green dots). **B, right panel** ABCA3 variants parsed by predicted effect. **C, left panel** 178 FBN1 variants are plotted by allele count and BayPR probabilities with disease assignment of P/LP. All 178 variants have probabilities >90%. **C, right panel** FBN1 (autosomal dominant Marfan syndrome) variants parsed by predicted effect. **D, left panel** 23 TGFBR1 (autosomal dominant Loeys-Dietz syndrome) variants assigned as P/LP are plotted by allele count and BayPR probabilities. **D, right panel** TGFBR1 variants parsed by predictive effect. **E, left panel** 54 TGFBR2 (autosomal dominant Loeys-Dietz syndrome) variants assigned as P/LP are plotted by allele count and BayPR probabilities. **E, right panel** TGFBR2 variants parsed by predicted effect. **F, left panel** 61 ABCD1 (X-linked adrenoleukodystrophy) variants previously assigned as P/LP or B/LB are plotted by allele count and BayPR probabilities. **F, right panel** ABCD1 variants parsed by predicted effect. **G, left panel** 143 TAZ (X-linked Barth syndrome) variants are plotted by allele count and BayPR probabilities, all with previous disease assignment of P/LP. **G, right panel** TAZ variants parsed by predicted effect.

We next tested the utility of BayPR for monogenic disorders with different Mendelian inheritance patterns. Marfan syndrome (caused by dysfunction of Fibrillin-1; *FBN1;* reported prevalence of 1:15,400 in Denmark)^22^ and Loeys-Dietz syndrome (caused by dysfunction of Transforming Growth Factor Beta 1 and 2; *TGFBR1* and *TGFBR2;* estimated prevalence <1;100,000)^23^ are autosomal dominant monogenic disorders. We applied the BayPR analysis to 178 *FBN1*, 23 *TGFBR1*, and 55 *TGFBR2* variants, all of which have been expertly curated as P/LP. All 178 FBN1 variants had probabilities >90% (**Figure 3C, left panel**), regardless of predicted effect (**Figure 3C, right panel**). The 23 P/LP variants in *TGFBR1* distributed such that two had probabilities >90%, 19 had 80-90%, while the remaining two were <10%, (**Figure 3D, left panel**). All variants were either missense (n=22) or in-frame deletion (n=1) (**Figure 3D, right panel**). All but 1 of 55 *TGFBR2* variants previously assigned as P/LP had probabilities >80% and 2 were >90% (**Figure 3E, left and right panels**). X-linked adrenoleukodystrophy (X-ALD) and X-linked Barth syndromes are monogenic disorders with estimated baseline prevalences of 1 in 20,000-50,000 and 1 in 300,000-400,000, respectively^24,25^. All 59 *ABCD1* (X-ALD) variants were assigned as P/LP by a single clinical DNA testing lab with extensive experience in interpretation of *ABCD1* variants^26^ and have probabilities >90%, while the one variant previously-assigned as B/LB has 0% probability. Parsing by predicted effect indicated all 52 missense variants have probabilities >90% (**Figure 3F, left and right panels**). We analyzed 143 variants in Tafazzin *(TAZ)* expertly curated as P/LP for Barth syndrome by the Human Tafazzin Gene Variants Database (https://www.barthsyndrome.org/research/tafazzindatabase.html). While only 41 (28.7%) *TAZ* variants showed probabilities >90%, 140 (97.9%) have probabilities >80%, and 3 (2.1%) have probabilities <10% (**Figure 3G, left panel**). All 63 missense *TAZ* variants have probabilities >80%, while the 2 of the P/LP variants with <10% probability are predicted to lead to complete loss-of-function (**Figure 3G, right panel**).

### Sensitivity and specificity of BayPR at different probability thresholds and different size datasets

We determined the sensitivity and specificity of BayPR at different thresholds on the three genes (*CFTR, ABCA3*, and *ABCD1*) that harbored P/LP and B/LB variants (**Table 1**). For this analysis, ‘true positives’ were variants annotated as P/LP that exceeded a probability of >90% or >80%; ‘true negatives’ were B/LB variants with probabilities <10% or <20%. Sensitivity of BayPR was greater than 95% for all three genes and for both combinations of probability. Likewise, specificity was greater than 94% for *CFTR* and *ABCD1* at 10% and 20% probabilities while specificity dramatically improved for *ABCA3* at the <20% threshold compared to the <10% threshold (100% vs. 60%). For the remaining disorders, an 80% threshold achieved correct assignment of over 97% of all known P/LP variants. To test how the size of the control dataset would affect BayPR probabilities, variants from all eight genes were re-analyzed using allele counts from a subset (genomes only; ∼15,000 individuals) of the full gnomAD dataset (exomes and genomes of ∼140,000 individuals). For *CFTR*, a large majority of P/LP variants moved from probabilities >90% in the full control dataset to probabilities between 80 and 90% in the genomes-only control dataset (**Table S1**). This reduction in probability was primarily driven by rare variants (**Figure S1**). In contrast, for *TAZ* and *TGFBR2*, 102 and 21 variants, respectively, moved from <90% probability in the full control dataset to >90% probability in the genomes-only dataset. For all other genes analyzed, variability in the control dataset size had little effect on probabilities and predictions of disease liability.

### BayPR can inform pathogenicity of variants variably associated with disease

To test if BayPR can differentiate underlying causes of variable disease association, we determined the probabilities of 41 variants of varying clinical consequence (VCC) (https://cftr2.org) and observed that 17 had predicted probabilities >80%, 16 had probabilities <20%, while the remaining 8 variants fall between these two thresholds (**Figure 4A**). Ten variants with probabilities <20% are reported to occur *in cis* with other *CFTR* variants. The low probabilities indicate these variants are likely benign when occurring alone but may contribute to pathogenicity in combination with other variants. Thus, variants of this type can occur in individuals with or without disease, thereby explaining their designation as VCC. *CFTR* variants designated as VCC with probabilities >80% could be benign or contributing to disease but in tight linkage disequilibrium with a pathogenic variant. We also applied BayPR to variants associated with mild hyperphenylalaninemia (MHP), which may or may not be clinically recognized. Of 58 MHP variants, 44 produced probabilities >80%, corroborating their disease association. Only 5 variants resulted in probabilities <20%, with the remaining 9 falling somewhere between (**Figure 4B**). These findings reveal population-based assessment can inform the interpretation of variants that can be variably associated with disease.

**Figure 4.**
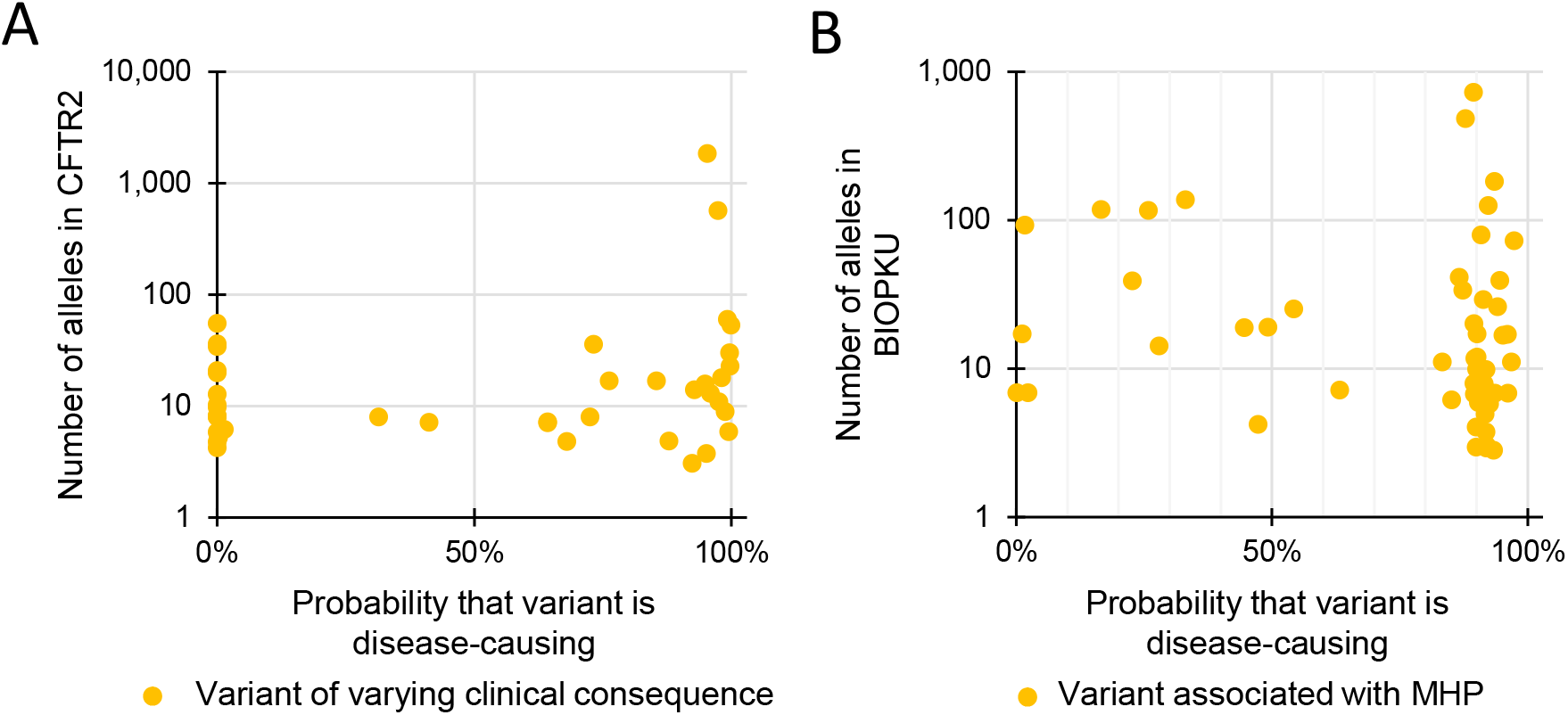
Variants associated with variable expressivity have a wide distribution of BayPR probabilities. Data is jittered for display purposes only in **Figures 4A-B**; precise allele counts and BayPR probabilities are provided in the Data Supplement. **A** 41 CFTR variants with disease assignment of varying clinical consequences (VCC) are plotted by BayPR probabilities and CFTR2 allele count and show a wide distribution. Sixteen variants have probabilities <20%, 17 variants have probabilities >80%, and eight are between 20 and 80%. **B** 58 PAH variants associated with mild hyperphenylaninemia (MHP) are plotted by BIOPKU allele count and BayPR probabilities and show a wide distribution. Forty-four have probabilities >80%, five variants have probabilities <10%, and the remaining nine variants fall between 20-80% probability of belonging to the disease-causing component.

## DISCUSSION

We demonstrate here that a component used to interpret variant pathogenicity has greater utility than previously recognized when deployed within a Bayesian framework. There are several distinct advantages to a Bayesian approach. First, probabilities can be generated for rare variants not previously observed in the general population. The rarity of such variants renders rigorous interpretation challenging, as it is difficult to confirm the variant’s existence in multiple unrelated individuals, obtain adequate clinical information for phenotypic evaluation and correlation and collect primary tissue samples or cells from a small number of affected individuals for functional assessment. Second, population-based approaches to variant interpretation, such as BayPR, are agnostic to variant mechanism and can reveal unexpected mechanisms of pathogenicity. For example, the c.371-2A>G variant of TAZ would be predicted to cause disease (alters +2 nucleotide of splice site) but BayPR generated a probability of <10%. Further evaluation revealed that the variant interrupts a splice site at the start of exon 5, an exon *excluded* in the prominent RNA isoform of TAZ in humans^27^. Thus, population data can provide an independent evaluation of variants that have functional testing results inconsistent with the associated phenotype and for variants not expected to affect the protein itself (e.g., synonymous changes), but are observed in affected individuals.

There are several potential limitations of variant interpretation methods based on population data alone. Within the control population, size matters; larger datasets should provide more accurate estimate of rare variants frequencies among presumed unaffected individuals. Our results partially support this supposition because BayPR prediction of *CFTR* variants (particularly for rare variants) improved when the full gnomAD data set (exomes and genomes) was used compared to genomes alone. Ongoing efforts to expand the size and diversity of ‘healthy’ population datasets should progressively improve the accuracy and utility of population-based approaches to variant interpretation^10,28,29^. A second consideration is phenotyping. Variants associated with mild presentation or late-onset disease could well result in some ‘affected’ individuals in the ‘healthy’ control population. Thus, the BayPR method will be less effective for late-onset conditions, unless appropriate age thresholds are applied to the control population. A third issue is applicability of prevalence ratios to *de novo* variants. Although our method is predicated on the distribution of transmitted alleles in healthy and affected individuals, BayPR generated accurate probabilities for the two X-linked disorders where *de novo* variants account for 40% (X-ALD)^30^ and 13% (Barth Syndrome)^14^ of cases. A fourth concern was over-counting P/LP variants due to the presence of related individuals (e.g. multiple affected siblings or affected relatives from different generations) in the ‘affected’ group. Over-counting, particularly of rare variants, can result in false assignment of a benign variant as disease-causing due to its biased prevalence ratio. In the U.S. CF population, there are 959 families with 1,962 affected twins or siblings among a sample of 30,000 affected individuals^31^, representing a potential over-counting of ∼3.3%. Over-counting could be more problematic in rarer disorders such as Barth Syndrome, due to diagnosis being more likely when two or more affected siblings are present, though curation of *TAZ* variants requires that variants only be submitted multiple times if they are observed in unrelated individuals (https://www.barthsyndrome.org/research/tafazzindatabase.html). However, the minor excess of alleles due to presence of siblings appeared to have no discernable decrease in the accuracy of BayPR for CF or for the rarer Mendelian disorders considered here (ILD and Barth syndrome). Finally, it is essential to consider the ancestry composition of the ‘affected’ and ‘healthy’ populations. Variants can be falsely assigned as disease-causing due to differences in the racial or ethnic makeup of the two samples^32^. Indeed, P/LP variants in *CFTR* and *PAH* occurring in samples well-represented in gnomAD (e.g., Latino, South Asian, or East Asian) but under-represented in the CFTR2 and BIOPKU datasets generated distorted prevalence ratios leading to low probabilities according to BayPR. Functional studies confirmed the variants are correctly assigned as P/LP. Therefore, it is recommended the effects of population stratification be considered prior to use of BayPR. Ideally, investigators should strive to achieve complete ascertainment of ‘affected’ subjects, which would lead to the capture of even very rare variants.

Population data provides a powerful tool for variant interpretation as recognized in the criteria developed by the ACMG/AMP and in the accuracy of interpretive algorithms incorporating population data^7,19^. Here we show a method based solely on variant counts can perform as well or better than popular interpretive tools based on multiple data sources and algorithms (including functional predictions). We interrogated a variety of other predictive tools^4,5,19,33-38^ and showed only ClinPred, which incorporates population-level data in a similar manner as our method, rivaled BayPR with regards to sensitivity and specificity. Important limitations to most of these tools include their utility for evaluation of missense variants only and inability to incorporate population data if a variant does not appear in a ‘healthy’ population dataset used as a comparison. In contrast, BayPR generates probabilities for even for ‘null’ variants that can provide independent evidence in situations where it is challenging to apply ‘PVS1’ criterion to assign pathogenicity (e.g. nonsense variants that do not elicit nonsense-mediated mRNA decay)^39^. The potential of population-based data utilized by BayPR emphasizes the importance of disease-specific datasets, such as CFTR2 and BIOPKU, that collect not only variants observed in ‘affected’ individuals, but also the *allele counts* of such variants, especially those suspected of being benign. While complete ascertainment of a disease population is ideal, we have demonstrated how multiple current locus-specific databases have sufficient data to enable accurate interpretation of variants by BayPR. Indeed, clinical and research labs could use their own variant counts for cases where there is reasonable confidence that individuals tested have the phenotype of interest to derive prevalence ratios and deploy BayPR to generate a probability score. While BayPR could be a standalone predictor of pathogenicity, it could be combined with orthogonal approaches such as functional testing or incorporated into guidelines such as those developed by the ACMG/AMP.

## DATA AND CODE AVAILABILITY

The datasets used for this study are available in the supplement. The code and associated documentation generated during this study are available at https://github.com/melishg/BayPR/.

## ACKNOWLEDGMENTS

This research would not have been possible without the generous contributions of patient and healthy subject data to the CFTR2 project, the BIOPKU and gnomAD datasets, and the Human *Tafazzin* Gene Variants Database.

## CONFLICT OF INTERESTS

The research was funded by the following organizations. The funders had no role in the conceptualization, design, data collection, analysis, decision to publish, or preparation of the manuscript.

JMC: NIH (R01 HL128475)

NB: This work is part of the RD-CONNECT initiative and was supported by a FP7-HEALTH-2012-INNOVATION-1 EU Grant (No. 305444).

LMN: NIH, American Thoracic Society, Eudowood Foundation

GRC: NIDDK (R01 DK44003), Cystic Fibrosis Foundation (CUTTIN13A1)

KSR, J Betz, TH, MAA, J Brown, HCD, GM, MBS, HJV, TAL: None

## ETHICS DECLARATION

All data were de-identified. The CFTR2 project is acknowledged by the Johns Hopkins University IRB (NA_00018599) and does not require IRB approval. All other datasets were obtained from publicly available resources and/or from researchers who provided de-identified variant information that meets NIH Exemption 4 (§46.104(d)(ii)) and does not qualify as human subjects research.

## AUTHOR CONTRIBUTIONS

Conceptualization: JMC, THB, TAL, and GRC; Data curation: KSR, NB, JBrown, HCD, GM, LMN, MBS, DJV, and GRC; Formal Analysis: JMC and JBetz; Funding acquisition: JMC, NB, LMN, and GRC; Investigation: JMC, KSR, JBetz, MAA, TAL, and GRC; Methodology: JBetz and TAL; Project administration: GRC; Resources: JBetz, KSR, NB, JBrown, HCD, GM, LMN, MBS, DJV, and GRC; Software: JBetz and TAL; Supervision: GRC; Validation: JMC, KSR, and GRC; Visualization: JMC, JBetz, and KSR; Writing – original draft: JMC, JBetz, KSR, and GRC; Writing – review & editing: JMC, JBetz, KSR, MAA, NB, JBrown, HCD, GM, LMN, MBS, DJV, THB, TAL, and GRC.

## WEB RESOURCES

https://cftr2.org

https://gnomad.broadinstitute.org/

http://www.biopku.org/home/home.asp

https://www.barthsyndrome.org/research/tafazzindatabase.html

https://useast.ensembl.org/Tools/VEP

