## Supplement for "Accurate assignment of disease liability to genetic variants using only population data"

**Assigning disease liability to genetic variants using prevalence ratios**

Supplementary Material

In addition to this file, supplementary data also includes one data supplement file, which contains all genetic data (control and disease populations) and disease liability assignments for alleles.

**Data Sources**

All data as described below were de-identified. The CFTR2 project is acknowledged by the Johns Hopkins IRB (NA_00018599) and does not require IRB approval. All other datasets were obtained from publicly available resources and/or from researchers who provided de-identified variant information that meets NIH Exemption 4 (§46.104(d)(ii)) and does not qualify as human subjects research. In some cases, variant names were provided to the analytic team in legacy terminology or protein name only. To obtain the associated nucleotide changes, original laboratory reports or records were reviewed by providing researchers when available. Additional HGVS cDNA and protein changes were obtained using VEP v10135.

**“Control” Variant:** **Data Source:** De-identified aggregated genomic data were downloaded from gnomAD v2.1.1, a publicly-available database consisting of 125,748 exome sequences and 15,708 whole genome sequences from unrelated individuals^27^. All genetic data (control and disease populations) and disease liability assignments for alleles can be found in the **data supplement**.

**Cystic Fibrosis (*CFTR*): Data Source:** For the CF disease population, de-identified *CFTR* genotype data in the CFTR2 [Clinical and Functional TRanslation of CFTR] (https://cftr2.org) database were provided for 89,052 individuals by national or regional CF patient registries or individual clinics from clinical records^19^. Individuals with no identified *CFTR* variants were excluded. Out of a total of 150,272 potential disease-causing alleles having received some genetic testing, 8,236 were not identified by available genetic testing and a total of 4,492 complex alleles, duplications/deletions greater than 3 base pairs, and variants in regions with poor or no coverage in gnomAD were excluded from analyses to yield a dataset of 137,544 *CFTR* allelic variants. **Curation:** Disease liability for specific alleles was determined by CFTR2 expert panels using clinical characteristics, functional studies, and penetrance analysis.

**Phenylketonuria (*PAH*): Data Source:** For the PKU disease population, de-identified *PAH* genotype data in the BIOPKU database were provided for 9,953 individuals^28,29^. Out of a total of 19,906 potential disease-causing alleles, 331 either were not identified by available genetic testing or were duplications/deletions greater than 3 base pairs and were excluded from analyses to yield a dataset of 19,575 *PAH* allelic variants. **Curation:** Disease liability of specific alleles was determined based on frequencies of the metabolic phenotype for genotypes presenting in a functionally hemizygous state and genotypes were compared to blood phenylalanine levels and BH4 responsiveness^20^.

**Interstitial Lung Disease (*ABCA3*):** **Data Source:** A list of de-identified variants was assembled from candidate gene sequencing was performed on symptomatic infants and children suspected of having genetic surfactant dysfunction in research laboratories at Washington University School of Medicine and Johns Hopkins University School of Medicine and critically reviewed publications^30^. **Data Curation:** Disease liability was determined by expert review of clinical characteristics, supporting imaging, family segregation studies, and/or lung histopathology if available, and mRNA splicing studies.

**X-linked Adrenoleukodystrophy (*ABCD1*): Data Source:** *ABCD1* variants were identified during clinical testing in the Johns Hopkins Genomics DNA Diagnostic lab from 2016 to 2020.  **Curation:** All variants were classified using current ACMG/AMP criteria^6^.

**Barth Syndrome (TAZ): Data Source:** De-identified *TAZ* variants were obtained from the Human Tafazzin Gene Variants Database (a sub-database of the Barth Syndrome Registry and Repository), which is publicly available through the Barth Syndrome Foundation (https://www.barthsyndrome.org/research/tafazzindatabase.html). Data were supplied by contributions from clinicians/families^31^. **Curation:** Disease liability was determined by expert review of clinical characteristics, cardiolipin/MLCL levels, and family segregation studies and/or mRNA splicing studies if available.

**Marfan Syndrome (*FBN1*): Data Source:** Allelic variants were identified during clinical testing at various CLIA-approved commercial labs from 2006 to 2020. **Curation:** Clinical data and variants are reviewed by a single clinical director and genetic counselor. All variants were classified using current ACMG/AMP criteria^6^.

**Loeys-Dietz Syndrome (*TGFBR1*/*TGFBR2*):** **Data Source:** Allelic variants were identified during clinical testing at various CLIA-approved commercial labs from 2006 to 2020. *SMAD3, TGFB2,* and *TGFB3* variants were not examined in this study. **Curation:** Clinical data and variants are reviewed by a single clinical director and genetic counselor. All variants were classified using current ACMG/AMP criteria^6^.

**Supplementary Methods**

### Summary

An empirical Bayesian approach was used to analyze the counts of allelic variants in the disease-specific population database relative to the control gnomAD population using a two component finite mixture model. In this empirical Bayes model, information across variants is pooled in order to get better estimates of each individual variant. Theoretically, this allows for more stable estimates, even when there may be few or no variants observed among the control cohort. The amount of information pooled is dependent on the number of variants observed, and the prior estimated from the data. With a richer dataset (the number of variants observed is larger, the total sample size is larger), there is less pooled information, but in a more restricted dataset (the number of variants observed is smaller, the total sample size is smaller), pooling increases. The algorithm is also dependent on the prior; when the prior is has a flatter distribution, reflecting more uncertainty about the distribution of prevalence ratios, the prior will exert less of an effect on the observed prevalence ratio. However, when the prior distribution is more peaked, reflecting less uncertainty about the distribution of prevalence ratios, it will exert more of an effect on the observed prevalence ratio.

Each component modeled the allele counts of specific allelic variants in the disease-specific database conditional on the total number of observations, using a binomial likelihood, placing a beta prior on a transformation of the prevalence ratio. Estimates were obtained by maximizing the marginal likelihood using the expectation-maximization (EM) algorithm implemented using the optimx package for R version (R Foundation for Statistical Computing, Vienna, Austria). A grid search was used to assess sensitivity to starting values for the EM algorithm, including the parameters of each mixture component and the mixing fraction. The model allows for variation in the size of each database (particularly for true for the control database used in this study), but does assume a constant ratio of the ratio of these database sizes; variation from this ratio would induce a different prior on the prevalence ratio. The R program used to generate the probabilities can be found on github (https://github.com/melishg/BayPR/).

For assessing the accuracy of our Bayesian algorithm, the probability of a variant belonging to one of the two beta distributions was compared to known disease liability obtained through functional and/or clinical data.

**Introduction**

For a particular genetic variant, denoted variant $k$, the binomial model allows us to infer about the proportion of variants arising from controls, denoted $\theta_{k}$. This model depends on $Y_{1k}$, the number of observed variants in cases among the $n_{1k}$ cases assessed for variant $k$, and $Y_{0k}$, the number of observed variants in controls among the $n_{0k}$ controls assessed for variant $k$. The total number of observed variants of type $k$ is denoted $T_{k}$, and the relative sample size in cases relative to controls for variant $k$ is denoted $r_{k}=(n_{1k}/n_{0k})$. We can transform the proportion of variants among the cases to get a prevalence ratio for variant $k$, denoted $\gamma_{k}$: this tells us how much more prevalent a variant is in cases relative to controls. The transformation from the proportion of variants arising from cases $(\theta_{k})$ to the prevalence ratio $(\gamma_{k})$ depends on the sample size ratio $r_{k}$.

If we have $K$ total variants, instead of viewing each variant in isolation, we can view them as a sample from a population of genetic variants. In this population of variants, the proportion of variants arising from controls follows a beta distribution, whose shape is determined by two parameters, $\alpha$ and $\beta$. When viewed in this way, each variant informs us about the distribution of the proportion of variants arising from controls in the larger population of variants, which helps provide more stable estimates for a particular variant when the observed frequency of that variant is low.

In this beta-binomial model, each variant is viewed arising from a population of variants, and in this population, the average proportion of variants arising from controls is $\mu=\alpha/(\alpha+\beta)$, and the variation in these proportions is $\mu(1-\mu)/(M+1)$, where $M=(\alpha+\beta)$. From a Bayesian viewpoint, the parameters of the beta distribution, $\alpha$ and $\beta$, can be interpreted as observing $\alpha$ prior variants among cases, and $\beta$ prior variants among controls, equivalent to data from a prior sample size of $M=(\alpha+\beta)$. This information can provide a stabilizing influence when the number of variants of a particular type are small or zero in one group. The stabilizing effect, called shrinkage, depends on the amount of observed data.

Let $s_{k}=M/(M+T_{k})$ denote the ratio of the prior sample size $M$ to the total sample size (prior sample size + number of variants of type $k$ observed). This $s_{k}$ can be thought of as a ‘shrinkage’ or ‘stabilization’ factor for variant $k$: when the number of observed variants is small relative to the prior sample size, the estimates are stabilized by being ‘pulled’ or ‘shrunken’ towards the population mean according to this shrinkage factor. As the number of observed variants becomes larger, this ratio will approach zero, and this shrinkage effect diminishes. We can also view this as a more principled, data-driven way of doing what some may tempted to do in practice: adding some small quantity to a numerator or denominator in order to avoid a numerical singularity.

Instead of viewing all variants as belonging to one homogenous population of variants, we could potentially view a sample variants as arising from a mixture of populations of variants. This is the conceptual basis of the mixture model. The added flexibility of this model allows for more local pooling of information across variants, and determining the sizes of population components and how compatible each variant is with each population component.

There are different methods for obtaining the parameters $\alpha$ and $\beta$, which describe the variation between variants in the proportion of variants arising from cases. Here, we used marginal maximum likelihood: finding the values of $\alpha$ and $\beta$ (equivalently $\mu$ and $M$) that maximize the marginal likelihood of the data, averaging over $\theta_{k}$. Since we are obtaining $\alpha$ and $\beta$ from the data itself, not from independent prior information, the ‘prior’ here can be thought of as a mechanism for stabilizing estimates that approximates a full Bayesian analysis and provides many desirable statistical properties.

### Outline

- 1. Why a Bayesian Approach?
- 2. The Probability Model
  - 2.1. The Binomial Likelihood
  - 2.2. The Beta-Binomial Model
  - 2.3. Bayes and Shrinkage
  - 2.4. The Beta Prime Distribution
  - 2.5. The Effect of $r_{k}$ on the Prior
- 3. Empirical Bayes Estimation
- 4. Using a Mixture Model

### 1. Why a Bayesian Approach?

Suppose we have information about the occurrence of $K$ genetic variants from a sample of cases from a population with a given disease and a sample of healthy controls from a population without that disease. One way of estimating the degree of association between a given genetic variant and disease status is to estimate the prevalence ratio: the ratio of genetic variant prevalence in cases relative to that in controls.

If we observe $Y_{1k}$ variants of type $k$ in $n_{1k}$ cases, and $Y_{0k}$ variants of type $k$ in $n_{0k}$ controls, we can estimate the prevalence in cases, denoted $\pi_{1k}$, using $\hat{\pi}_{1k}=Y_{1k}/n_{1k}$, and the prevalence in controls, denoted $\pi_{0k}$, using $\hat{\pi}_{0k}=Y_{0k}/n_{0k}$. We can estimate the prevalence ratio in cases relative controls, $\gamma_{k}$, using the ratio of the prevalence estimates: $\hat{\gamma}_{k}=(\hat{\pi}_{1k}/\hat{\pi}_{0k})=(Y_{1k}/n_{1k})/(Y_{0k}/n_{0k})$. Since it is the ratio of two probabilities, the prevalence ratio could take on any positive value: ratios greater than 1 indicate higher prevalence among cases, and ratios less than 1 indicate higher prevalence among controls.

One issue that may arise is that variants may be extremely rare in one population or another, which may result in unstable prevalence ratio estimates. If few or no variants are observed among controls, the denominator of the ratio becomes very small, resulting in unstable estimates of the prevalence ratio. Bayesian models augment the observed data with a prior distribution, which can provide estimates that are more stable and also have good frequentist statistical properties, such as interval estimate coverage and mean squared error.

### 2. The Probability Model

#### 2.1 The Binomial Likelihood

For each genetic variant $k=1, 2, \ldots, K$, we assume that we have a random sample of $n_{1k}$ cases and $n_{0k}$ controls who were evaluated for that variant, with $Y_{1k}$ occurrences in cases and $Y_{0k}$ occurrences in controls. The prevalence of variant $k$ in cases is denoted by $\pi_{1k}$, and the prevalence in controls is denoted by $\pi_{0k}$. The proportion of individuals in a sample of size $n$ of a variant $k$ independently sampled from a population with prevalence $\pi_{k}$ can be modeled using a binomial distribution:

$$Y\sim Bin(n,\pi): Pr\{Y=y|n,\pi\}=\binom{n}{y}\pi^{y}(1-\pi)^{n-y}$$

This binomial likelihood can be approximated using a Poisson distribution with rate parameter $\lambda=n\pi$ if $n$, the number of observations, is large and $\pi_{k}$, the prevalence of the variant, is small.

$$Y\sim Poisson(\lambda=n\pi): Pr\{Y=y|n,\pi\}=\frac{(n\pi)^{y}e^{-n\pi}}{y!}$$

If we observe $Y_{1k}$ variants of type $k$ in cases, $Y_{0k}$ variants of type $k$ in controls, let $T_{k}$ denote their sum. We can model the proportion of type $k$ variants that arose from cases out of the the total number of type $k$ variants observed, denoted $\theta_{k}$, using a binomial distribution:

$$Y_{0k}\sim Poisson(n_{0k}\pi_{0k}), Y_{1k}\sim Poisson(n_{1k}\pi_{1k}); Y_{0k}\perp Y_{1k}: \left( Y_{1k}|Y_{0k}+Y_{1k}=T_{k} \right)\sim Bin(T_{k},\theta_{k}),$$

where $\theta_{k}=(n_{1k}\pi_{1k})/(n_{1k}\pi_{1k}+n_{0k}\pi_{0k})$, the rate of occurrences in cases divided by the sum of the rates in cases and controls. We can divide the numerator and denominator by $\pi_{0k}$ to parameterize this distribution by the prevalence ratio $\gamma_{k}$:

$$\theta_{k}=\frac{n_{1k}\pi_{1k}/\pi_{0k}}{n_{1k}\pi_{1k}/\pi_{1k}+n_{0k}\pi_{0k}/\pi_{1k}}=\frac{n_{1k}\gamma_{k}}{n_{1k}\gamma_{k}+n_{0k}}$$

Let $r_{k}=n_{1k}/n_{0k}$ denote the ratio of the sample size of cases relative to controls. Dividing the numerator and denominator again by $n_{0k}$, the control sample size, gives:

$$\theta_{k}=\frac{n_{1k}\gamma_{k}/n_{0k}}{n_{1k}\gamma_{k}/n_{0k}+n_{0k}/n_{0k}}=\frac{r_{k}\gamma_{k}}{r_{k}\gamma_{k}+1}$$

Parameterizing the model this way allows the modeling multiple variants, allowing for variation in sample sizes across different variants, as long as their ratio of sample sizes for any given variant $r_{k}$ is comparable.

#### 2.2 The Beta-Binomial Model

Bayesian statistics augments the likelihood probability model with a prior probability model. Our model for the data is a binomial distribution, which depends on $n=T_{k}$, the number of variants observed and $\pi=\theta_{k}$, the proportion of variants arising from the population of cases. Instead of viewing each of our $K$ variants separately, we could view them as a sample from a population of genetic variants, and the proportions of variants due to cases in this population follow a probability distribution called the Beta distribution.

The shape of the beta distribution is controlled by two parameters, $\alpha$ and $\beta$. Depending on the values these parameters, the beta distribution can take on many possible shapes, as seen in the figures below. The mean of the beta distribution is given by $E[\theta_{k}|\alpha,\beta]=\mu=\alpha/(\alpha+\beta)$, and its variance is given by $var[\theta_{k}|\alpha,\beta]=\alpha\beta/\left( (\alpha+\beta)^{2}(\alpha+\beta+1) \right)$.

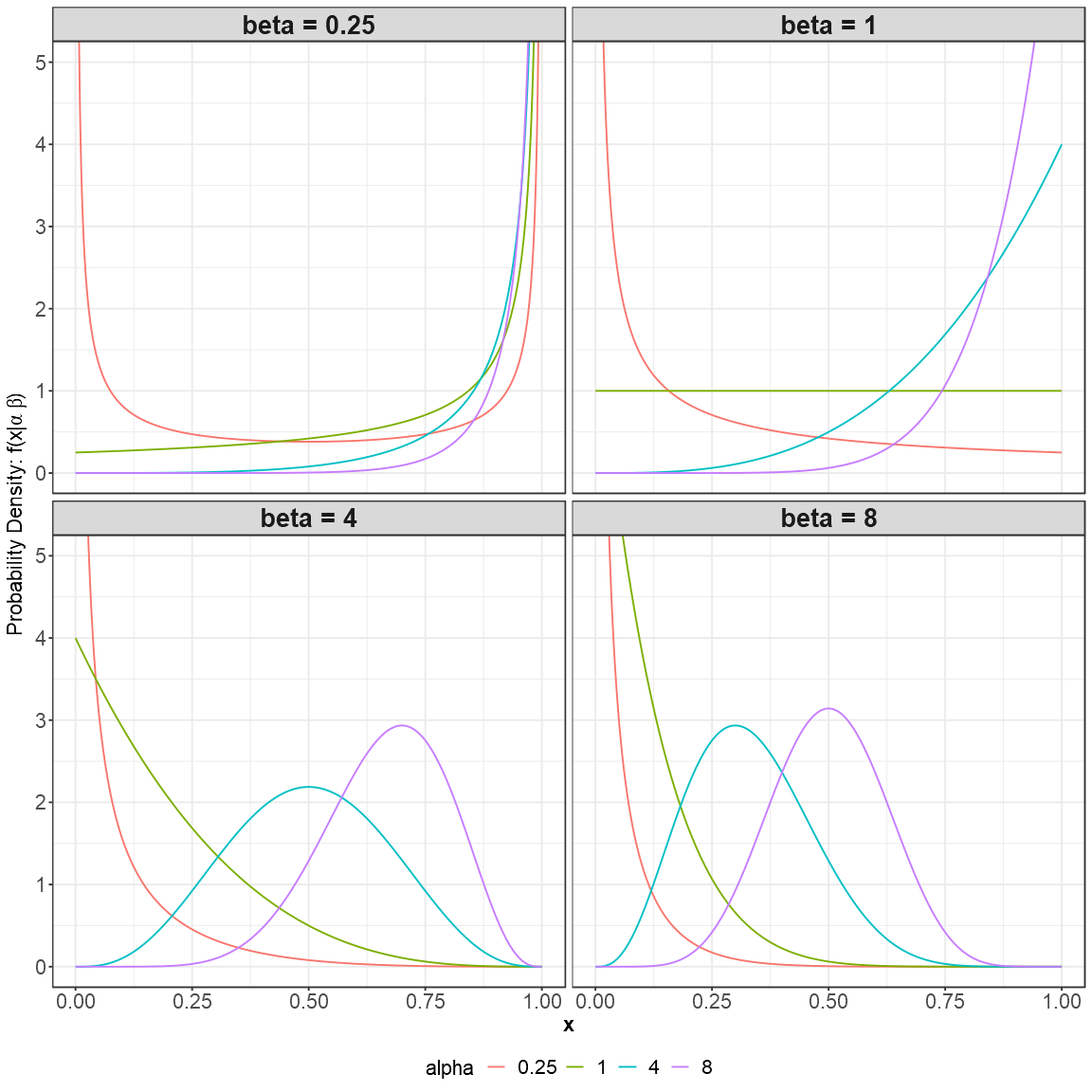

**Supplementary Figure 1:** Examples of beta distributions obtained by varying the values of the parameters alpha, indicated by the color, and beta, indicated by the figure panel.

We can also re-parameterize the beta distribution in terms of $\mu$, its mean, and $M=\alpha+\beta$:

$$Pr\{\theta_{k}|\mu,M\}=\frac{\Gamma(M)}{\Gamma(\mu M)\Gamma((1-\mu)M)}\theta^{\mu M-1}(1-\theta)^{(1-\mu)M-1}$$

In this parameterization, $\alpha=\mu M$ and $\beta=(1-\mu)M$. We can think of $M$ as a prior sample size: $\alpha$ prior variants among cases and $\beta$ prior variants among controls. When parameterized this way, the variance of the beta distribution is given by $var[\theta_{k}|\mu, M]=\mu(1-\mu)/(M+1)$.

The combination of a beta distribution for the proportion of variants arising from cases, and the binomial likelihood for the number of variants observed in cases, gives a Beta-Binomial Distribution. The parameters of the beta distribution are called hyperparameters, because their values determine the distribution of the proportion parameter in the binomial model.

The probability density function for the prior is:

$$Pr\{\theta_{k}|\alpha, \beta\}=\frac{\Gamma(\alpha+\beta)}{\Gamma(\alpha)\Gamma(\beta)}\theta_{k}^{\alpha-1}(1-\theta_{k})^{\beta-1}$$

The likelihood for the observed data is:

$$Pr\{Y_{1k}|T_{1k}, \theta_{k}\}=\binom{T_{1k}}{Y_{1k}}\theta_{k}^{Y_{1k}}(1-\theta_{k})^{Y_{1k}-T_{1k}}$$

Note the similarity between these two functions: both are products of the terms $\theta_{k}$ and $(1-\theta_{k})$. When combined using Bayes’ Rule, they give a posterior distribution is also a beta distribution:

$$Pr\{\theta_{k}|Y_{1k}, T_{k}, \alpha, \beta\}=\frac{\Gamma(\alpha+\beta+T_{k})}{\Gamma(\alpha+Y_{1k})\Gamma(\beta+Y_{0k})}\theta^{\alpha+Y_{1k}-1}(1-\theta)^{\beta+Y_{0k}-1}$$

This is just a beta distribution with $\alpha'=\alpha+Y_{1k}$ and $\beta'=\beta+Y_{0k}$: this is why we can interpret $\alpha$ as the prior number of variants observed in cases, and $\beta$ as the prior number of variants observed among controls. The posterior distribution, and its summaries (such as its mean, variance, and quantiles), allow us to infer about $\theta_{k}$, the proportion of variants due to cases, for each of our $K$ variants.

#### 2.3 Bayes and Shrinkage

The advantage of the Bayesian approach is that the prior distribution acts as a stabilizing influence when the number of observed variants is small, and this influence diminishes as the number of observed variants becomes larger. The effect of the prior can be more easily understood when viewed through the parameterization of $\mu$ and $M$. The mean of the posterior distribution is given by $E[\theta_{k}|\alpha,\beta]=\mu=\alpha/(\alpha+\beta)$. Re-writing this in terms of $\mu$ and $M$ gives:

$$E[\theta_{k}|\alpha,\beta]=\frac{\alpha+Y_{1k}}{\alpha+\beta+T_{k}}=\frac{\mu M+Y_{1k}}{M+T_{k}}=\frac{M}{T_{k}+M}\mu+\frac{T_{k}}{T_{k}+M}\left( \frac{Y_{1k}}{T_{k}} \right)=s_{k}\mu+(1-s_{k})\left( \frac{Y_{1k}}{T_{k}} \right)$$

Since $M$ acts as a prior sample size, and $T_{k}$ is the total number of variants observed, the quantity $s_{k}=M/(M+T_{K})$ represents the proportion of the information coming from the prior, and $(1-s_{k})=T_{k}/(M+T_{K})$ represents the proportion of information coming from the observed data. Here we see the posterior mean is a combination of the prior mean $\mu$ and the proportion of variants among cases to the total number of variants observed $(Y_{1k}/T_{k})$, each weighted according their sample size contribution. When $T_{k}$, the number of observed variants is small, the posterior mean is ‘pulled’ or ‘shrunken’ towards the prior mean: this gives more stable estimates. As the number of observed variants becomes increasingly larger than the prior sample size, the effect of the prior diminishes.

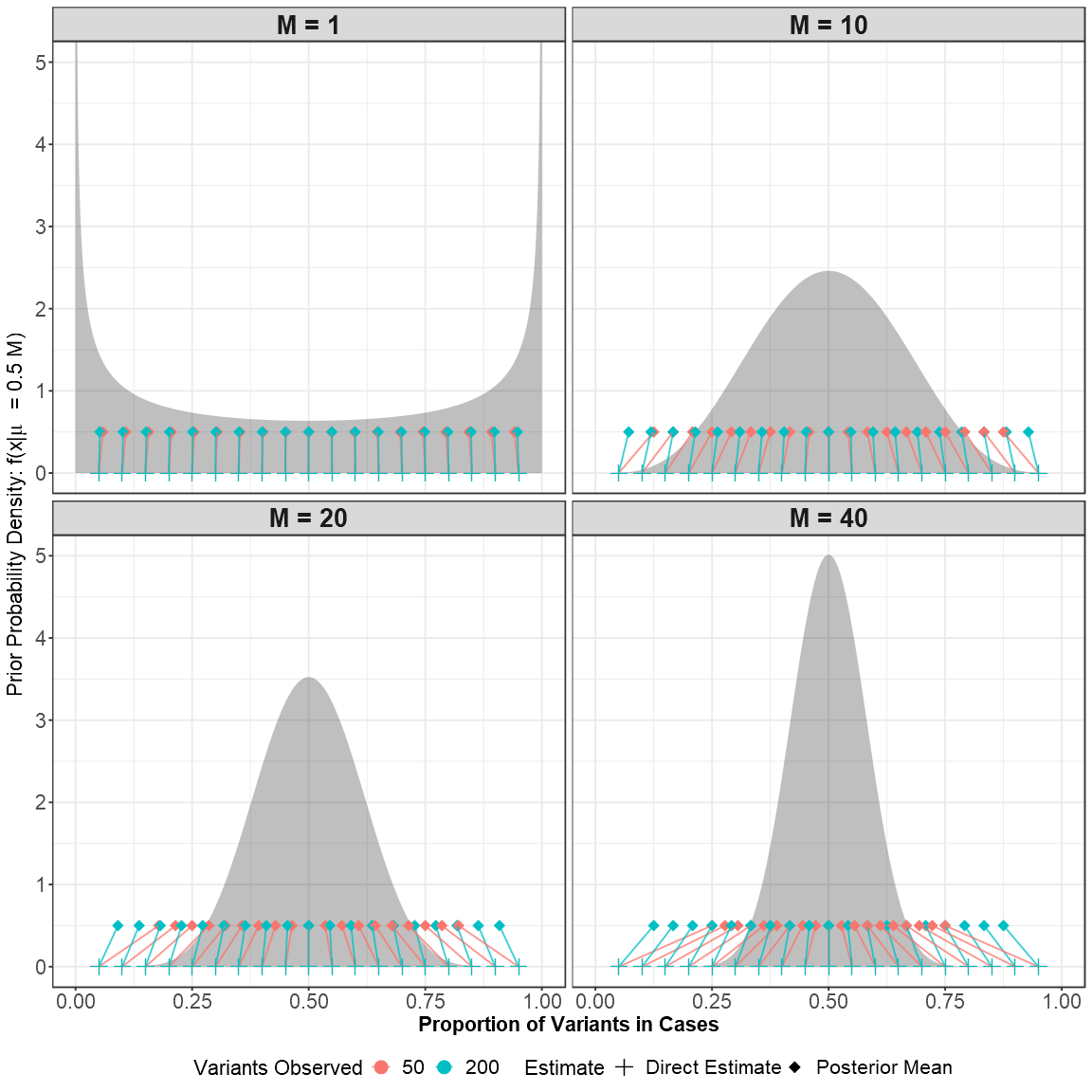

**Supplementary Figure 2:** An example of how the shape of the prior affects the posterior mean, and how this depends on both the number of observed variants, the proportion of variants seen in cases, and the prior sample size. The prior distribution is shown in gray. When the prior sample size (M) is small, the effect of the prior is smallest, and the posterior means are very close to the direct estimates. As the prior sample size increases, direct estimates are ‘pulled’ or ‘shrunken’ towards the prior mean (here, 0.5), with greater shrinkage occurring when fewer variants are observed.

#### 2.4 The Beta Prime Distribution

The beta distribution describes the distribution of a random variable over the interval $(0,1)$, which is convenient for describing a proportion or probability. However, if we want to infer about the prevalence ratio $\gamma_{k}$, we need to transform back to the prevalence ratio scale:

$$\frac{\theta_{k}}{r_{k}(1-\theta_{k})}=\gamma_{k}$$

Notice that when $r_{k}=1$ (sample sizes are identical between cases and controls), this transformation is from the probability scale to the odds scale. If we have a beta random variable, and transform it to the odds scale, this results in a variable with the beta prime distribution. The shape of this distribution is governed by the same parameters, $\alpha$ and $\beta$: If $X\sim Beta(\alpha, \beta)$ then $Y=X/(1-X)\sim Beta Prime(\alpha, \beta)$.

If $(Y|\alpha,\beta)\sim Beta Prime(\alpha,\beta)$, its mean is given by $E[Y|\alpha,\beta]=\alpha/(\beta-1)$, and its variance is given by $var[Y|\alpha,\beta]=\alpha(\alpha+\beta-1)/\left( (\beta-2)(\beta-1)^{2} \right)$

For a beta-prime posterior distribution of prevalence ratios, we can plug in our posterior values of $\alpha'=\alpha+Y_{1k}$ and $\beta'=\beta+Y_{0k}$. When $r_{k}\neq1$ (the sample size differs between cases and controls), the posterior becomes a scaled version of the beta prime distribution. The mean is scaled by $1/r_{k}$, and the variance is scaled by $1/r_{k}^{2}$:

$$E[\gamma_{k}|Y_{1k},T_{k},\alpha,\beta]=\frac{1}{r_{k}}\frac{\alpha+Y_{1k}}{(\beta+Y_{0k}-1)}=\frac{\frac{\alpha}{n_{1k}}+\frac{Y_{1k}}{n_{1k}}}{\frac{\beta-1}{n_{0k}}+\frac{Y_{0k}}{n_{0k}}}$$

By adding $\alpha/n_{1k}$ to the numerator, and $(\beta-1)/n_{0k}$ to the denominator, this stabilizes the prevalence ratio. If we think of $\alpha$ as the prior number of variants observed among cases, and $\beta$ as the prior number of variants among controls, the numerator is ‘pulled’ or ‘shrunken’ towards $\alpha$, as the denominator is towards $\beta$. This pull becomes negligable as the sample size becomes large.

This can also be seen when we paramaterize instead using $\mu$ and $M$:

$$E[\gamma_{k}|Y_{1k},T_{k},\mu,M]=\frac{\frac{\mu M}{n_{1k}}+\frac{Y_{1k}}{n_{1k}}}{\frac{(1-\mu)M-1}{n_{0k}}+\frac{Y_{0k}}{n_{0k}}}$$

The numerator is ‘pulled’ or ‘shrunken’ towards the prior mean $\mu$, and this effect diminishes as $n_{1k}$, the sample size among cases, becomes large compared to the prior sample size $M$. The denominator is ‘pulled’ or ‘shrunken’ towards $(1-\mu)$, and this effect diminishes as $n_{0k}$, the sample size among controls, becomes large compared to the prior sample size $M$.

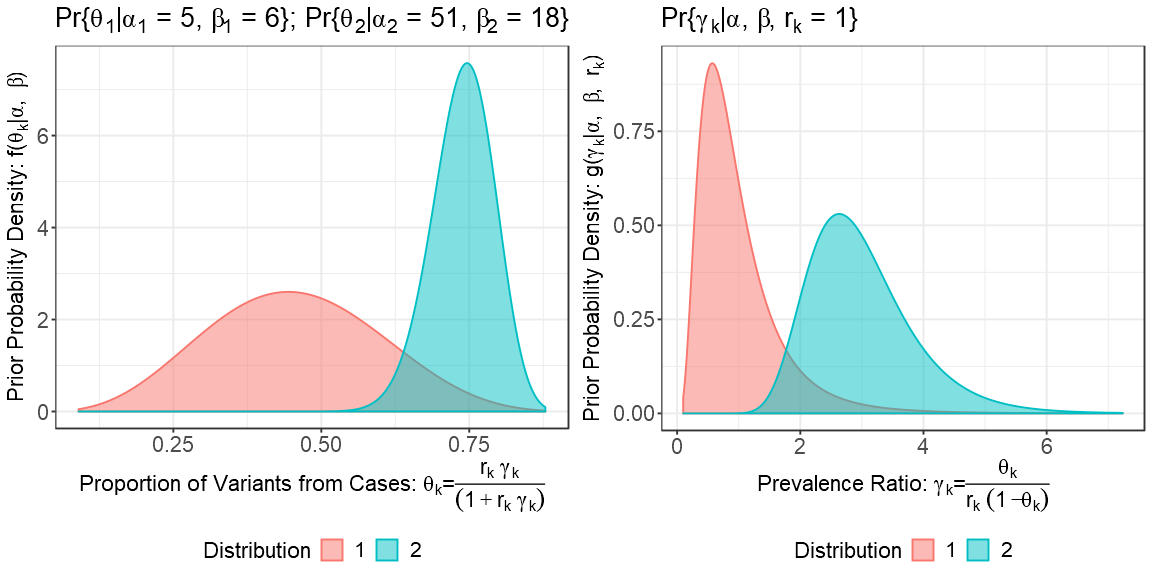

**Supplementary Figure 3:** Examples of two beta distributions, modeling the proportion of variants observed in cases, and their corresponding distributions of prevalence ratios.

#### 2.5 The Effect of $\boldsymbol{r}_{\boldsymbol{k}}$ on the Prior

Note that the transformation to the prevalence ratio scale depends on $r_{k}$, the ratio of sample sizes in cases $(n_{1})$ relative to controls $(n_{0})$. Note how the same beta model could represent populations of variants with lower, equal, or higher prevalence in cases relative to controls, depending on the value of $r_{k}:$

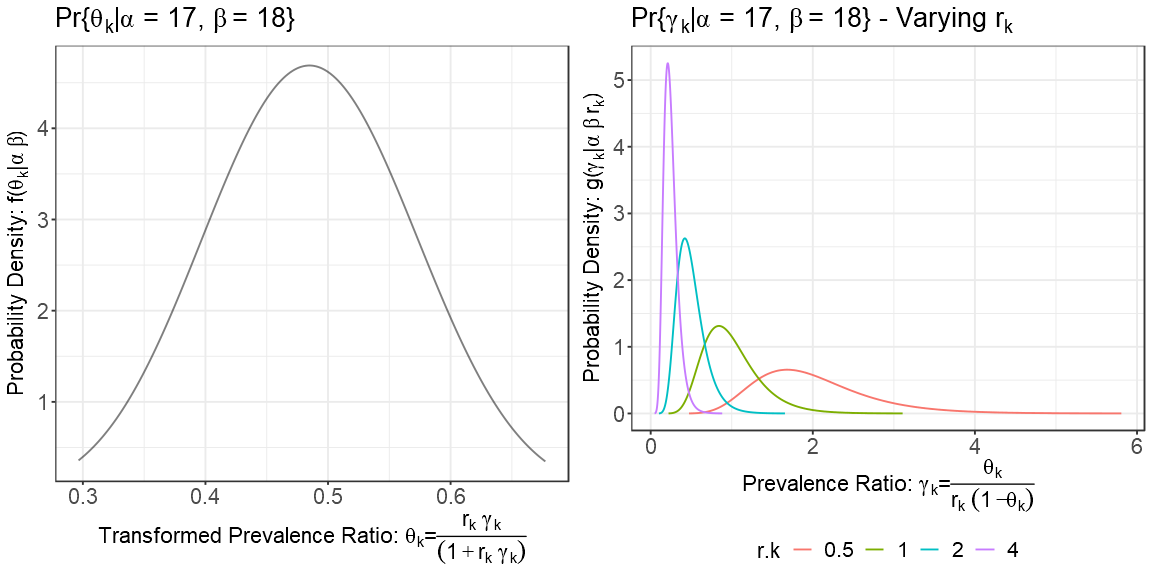

**Supplementary Figure 4:** Example of how one beta distribution, could represent lower, equal, or higher prevalence between cases and controls, depending on the relative sample size between cases and controls.

This also illustrates the importance of the assumption about the variation $r_{k}$ across all $K$ variants: if there is appreciable variation in this ratio across variants, then variants will essentially be ‘pulled’ or ‘shrunken’ in different directions.

### 3. Empirical Bayes Estimation

Up until now, we have been treating the parameters of our beta prior distribution, $\alpha$ and $\beta$, as known quantities, and showing how their values ‘pull’ or ‘shrink’ the direct estimates towards the prior mean on on the proportion (or $\theta_{k}$) scale, and ‘pull’ or ‘shrink’ the numerator and denominator on the ratio scale. But how do we determine suitable values for these ‘hyperparameters?’

Rather than either supplying exact values for these parameters, or specifying a prior distribution over these parameters, we can find the values of these parameters that maximize the *marginal likelihood* of the observed data, averaging over the parameter $\theta_{k}$: $Pr\{Y_{1k}|T_{k}, \mu, M\}$

$$Pr\{Y_{1k}|T_{k}, \mu, M\}=f(Y_{1k}|T_{k}, \mu, M)=\frac{\Gamma(T_{k}+1)}{\Gamma\left( Y_{1k}+1 \right)\Gamma\left( T_{k}-Y_{1k}+1 \right)}\frac{\Gamma(Y_{1k}+\mu M)\Gamma\left( T_{k}-Y_{1k}+(1-\mu)M \right)}{\Gamma\left( T_{k}+M \right)}\frac{\Gamma\left( M \right)}{\Gamma(\mu M)\Gamma\left( (1-\mu)M \right)}$$

where $\Gamma$ denotes the gamma function.

### 4. Using a Mixture Model

Instead of assuming that all variants are represented by one beta distribution that describes the proportion of variants among cases, we can relax this assumption by assuming that the observed data were generated from a mixture of different beta distributions.

Instead of our model having only two parameters, $\mu$ and $M$, our mixture model will have 5 parameters: $\mu_{1}$ and $M_{1}$, which control the shape of one beta distribution, $\mu_{2}$ and $M_{2}$, which control the shape of the second beta distribution, and $\epsilon$, which is the proportion of variants arising from the second beta distribution:

$$f(Y_{1k}|T_{k}, \mu_{1}, M_{1}, \mu_{2}, M_{2}, \epsilon)=\epsilon f(Y_{1k}|T_{k}, \mu_{1}, M_{1})+(1-\epsilon)f(Y_{1k}|T_{k}, \mu_{2}, M_{2})$$

Note that two different parameter vectors $(\mu_{1}=a, M_{1}=b, \mu_{2}=c, M_{2}=d,\epsilon=e)$ and $(\mu_{1}=c, M_{1}=d, \mu_{2}=a, M_{2}=b,\epsilon=1-e)$ result in identical values of the mixture model: for this reason, the constraint $\epsilon<0.5$ is imposed to identify a unique solution. Estimates of these parameters are obtained by maximizing the marginal likelihood of the mixture.

Instead of each direct estimate being ‘pulled’ or ‘shrunken’ in the same direction, each distribution will exert a different ‘pull’ on the data, with the ‘pull’ being related to the relative compatibility between each model component and the data.

From this mixture distribution, in addition to obtaining posterior means, variances, and quantiles, we can additionally obtain:

- The marginal likelihood ratio: $MLR_{2/1}=Pr\{Y_{1k}|T_{k}, \mu_{2}, M_{2}\}/Pr\{Y_{1k}|T_{k}, \mu_{1}, M_{1}\}$
- The posterior odds of belonging to component 2 vs. 1: $PO_{2k}=(\epsilon/(1-\epsilon))MLR_{2/1}$
- The posterior probability of belonging to component 2: $P_{2k}=PO_{2k}/(1+PO_{2k})$

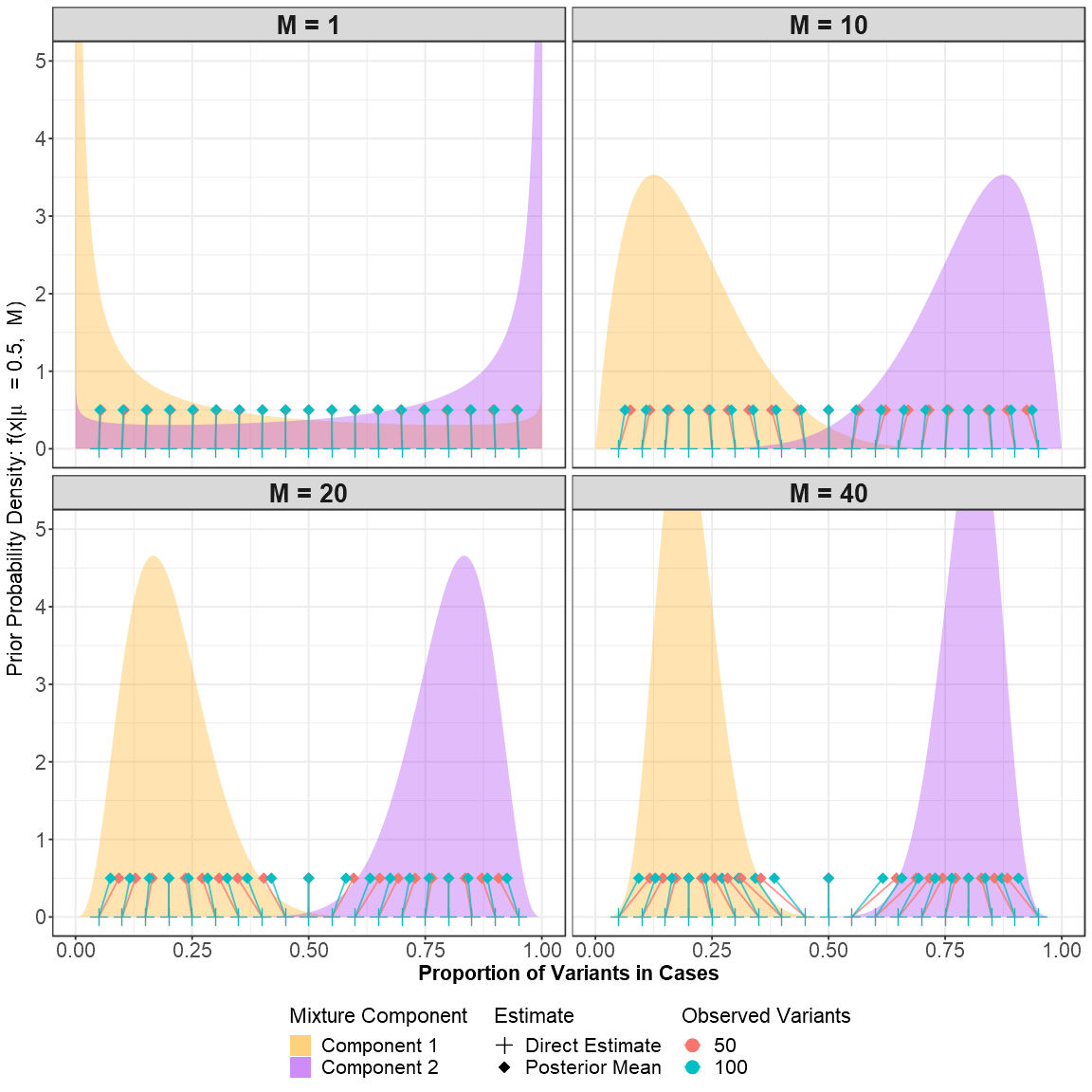

**Supplementary Figure 5:** An example of a mixture prior, and the effect of each mixture component on the direct estimate. When the prior sample size (M) is small, the effect of the prior is smallest, and the posterior means are very close to the direct estimates. As the prior sample size increases, direct estimates are ‘pulled’ or ‘shrunken’ towards each of the priors. The ‘pull’ of each prior depends on the marginal likelihood ratio: the relative compatibility between the direct estimate and each prior probability component.

**Supplementary Results**

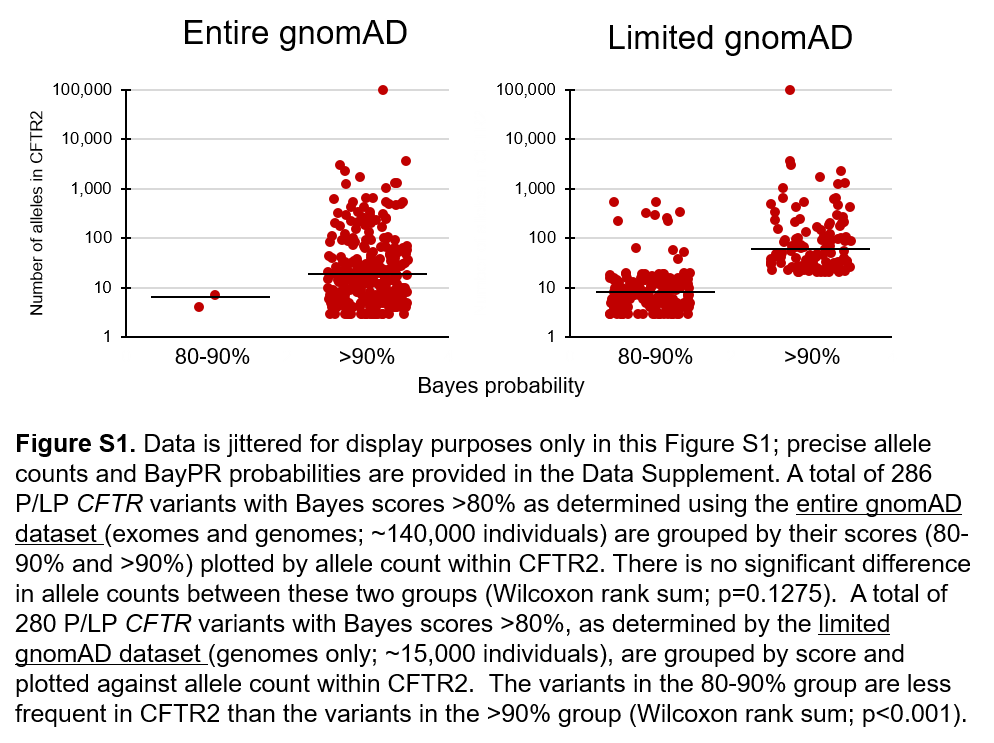

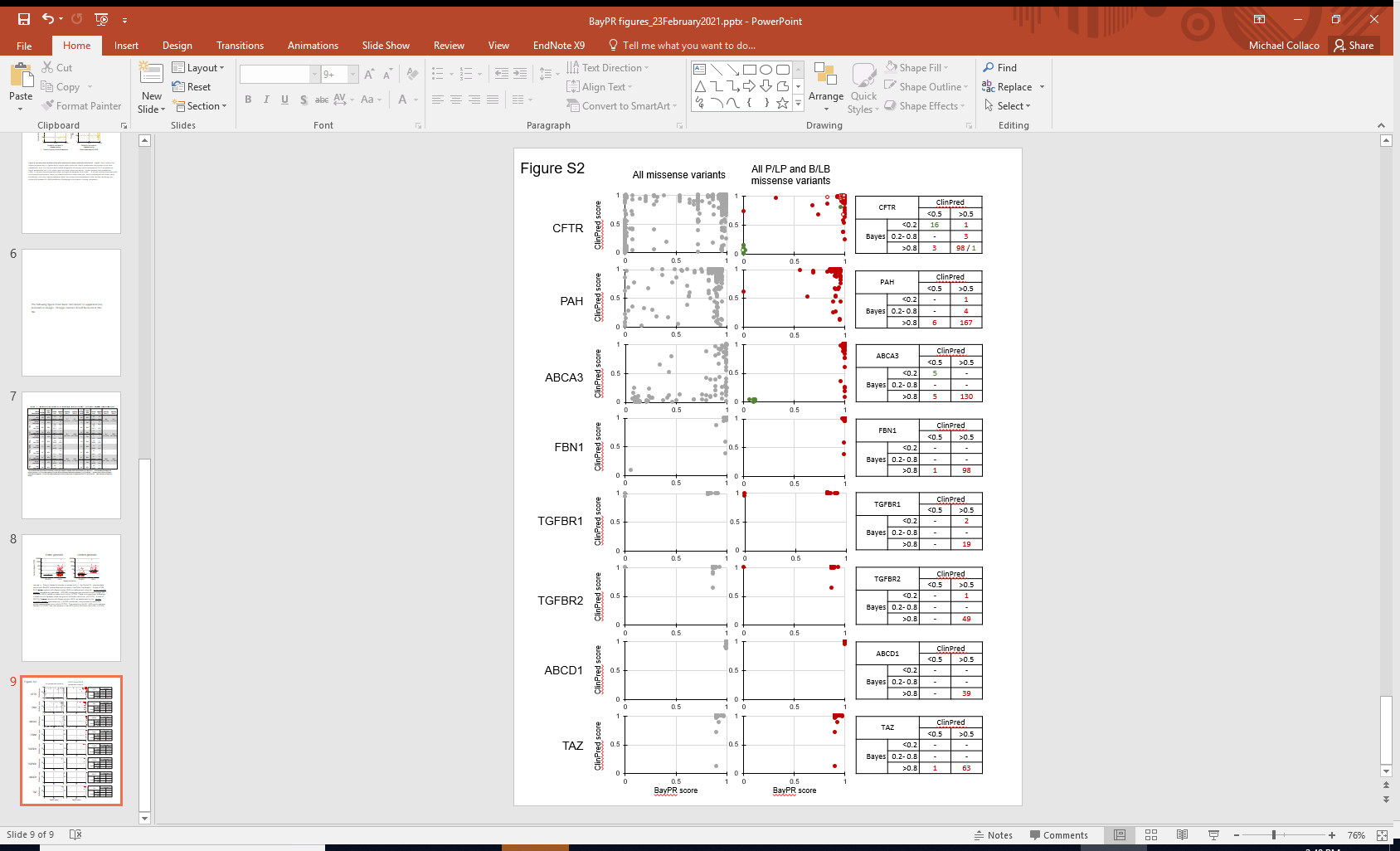

**Figure S2. Legend:** Expertly-assigned variants associated with recessive, dominant, and X-linked conditions plotted to compare their BayPR and ClinPred scores. The open circles shown for CFTR (both red and green) indicate variants newly-assigned as P/LP or B/LB, respectively (see **Figure 2,** panels C and D). Tables indicate distribution of variant scores across disease liability thresholds using BayPR and ClinPred.

**Table S1. Sensitivity and specificity analysis using a limited gnomAD dataset (genomes only)**

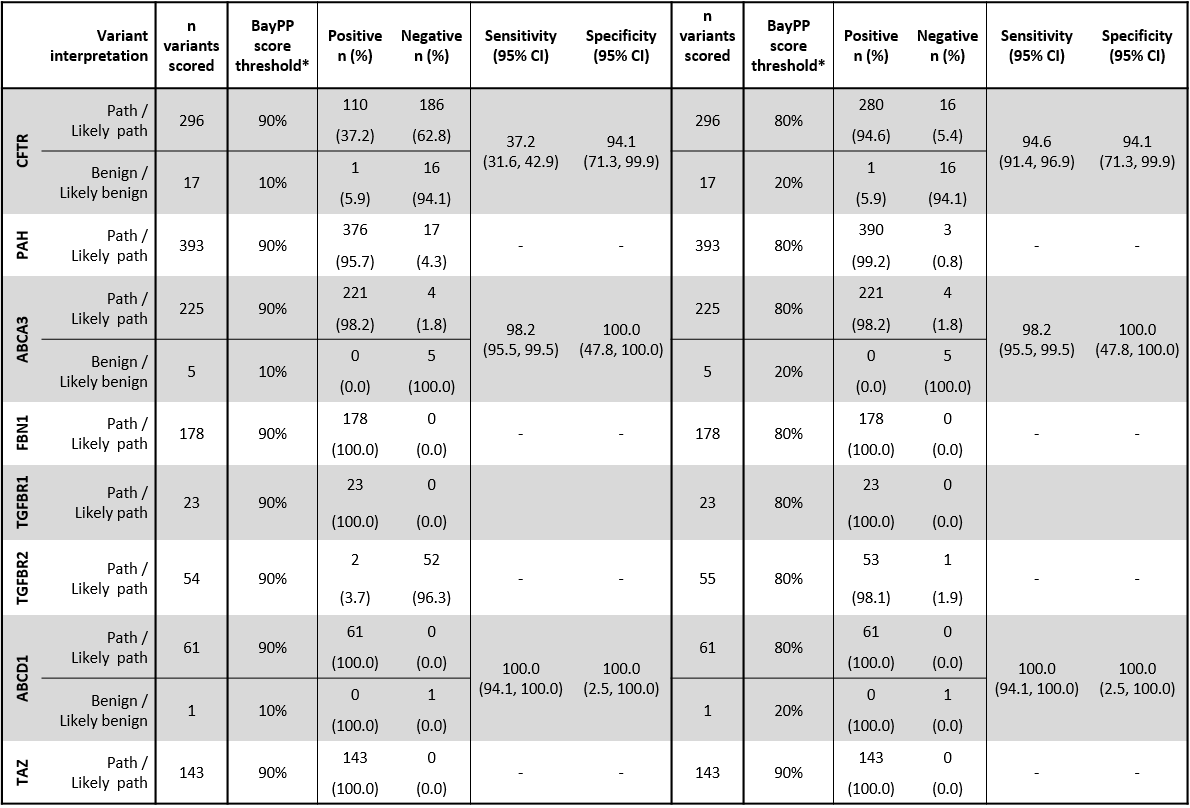

*Score thresholds were set to determine whether a variant had a positive or negative result from the Bayesian XYZ algorithm for the purposes of sensitivity and specificity analysis. For P/LP variants, Bayesian XYZ scores >90% or >80% were considered true positives. For B/LB variants, Bayesian scores <10% or <20% were considered true negatives. P/LP and B/LB determinations were made by expert review. Only genes with both P/LP and B/LB underwent sensitivity and specificity analysis.

*Score thresholds were set to determine whether a variant had a positive or negative result from the Bayesian XYZ algorithm for the purposes of sensitivity and specificity analysis. For P/LP variants, Bayesian XYZ scores >90% or >80% were considered true positives. For B/LB variants, Bayesian scores <10% or <20% were considered true negatives. P/LP and B/LB determinations were made by expert review. Only genes with both P/LP and B/LB underwent sensitivity and specificity analysis.
